# Shedding light on the contribution of amygdalar excitatory projections to prepulse inhibition of the auditory startle reflex

**DOI:** 10.1101/2021.01.24.427973

**Authors:** Jose Carlos Cano, Wanyun Huang, Karine Fénelon

## Abstract

**Background:** Sensorimotor gating is a fundamental neural filtering process that allows attention to be focused on a given stimulus. Sensory gating, commonly measured using the prepulse inhibition (PPI) of the auditory startle reflex task, is impaired in patients suffering from various neurological and psychiatric disorders, including schizophrenia. Because PPI deficits are often associated with attention and cognitive impairments, they are widely used as biomarkers in pre-clinical research for anti-psychotic drug screening. Yet, the neurotransmitter systems and synaptic mechanisms underlying PPI are still not resolved, even under physiological conditions. Recent evidence ruled out the longstanding hypothesis that PPI is mediated by midbrain cholinergic inputs to the caudal pontine reticular nucleus (PnC). Instead, glutamatergic, glycinergic and GABAergic inhibitory mechanisms are now suggested to be crucial for PPI, at the PnC level. Since amygdalar dysfunctions affect PPI and are common to pathologies displaying sensorimotor gating deficits, the present study was designed to test that direct projections to the PnC originating from the amygdala, contribute to PPI.

**Results:** Using Wild Type and transgenic mice expressing eGFP under the control of the glycine transporter type 2 promoter (GlyT2-eGFP mice), we first employed tract-tracing, morphological reconstructions and immunohistochemical analyses to demonstrate that the central nucleus of the amygdala (CeA) sends glutamatergic inputs latero-ventrally to PnC neurons, including GlyT2^+^ cells. Then, we showed the contribution of the CeA-PnC excitatory synapses to PPI *in vivo* by demonstrating that optogenetic inhibition of this connection decreases PPI, and optogenetic activation induces partial PPI. Finally, in GlyT2-Cre mice, whole-cell recordings of GlyT2^+^ PnC neurons *in vitro* paired with optogenetic stimulation of CeA fibers, as well as photo-inhibition of GlyT2^+^ PnC neurons *in vivo*, allowed us to implicate GlyT2^+^ neurons in the PPI pathway.

**Conclusions:** Our results uncover a feed-forward inhibitory mechanism within the brainstem startle circuit by which amygdalar glutamatergic inputs and GlyT2^+^ PnC neurons play a key role in meditating PPI. We are providing new insights to the clinically-relevant theoretical construct of PPI, which is disrupted in various neuropsychiatric and neurodevelopmental diseases.

## Background

Sensorimotor gating is the ability of a weak sensory event to inhibit or “gate” the motor response to an intense sensory stimulus. Such as fundamental neuronal process is thought to help attention to be oriented towards a given stimulus. Currently, prepulse inhibition (PPI) of the acoustic startle reflex task remains the gold standard operational measure of sensorimotor gating [1–3], used both in humans and various animal models [2, 4]. PPI is a paradigm in which a pre-stimulus of low intensity (“prepulse”) presented ∼10-500ms before a startle stimulus (“pulse”), attenuates the startle response [1–9]. PPI suppresses other, potentially interfering sensory or behavioral responses, to ensure efficient information processing. Abnormal PPI arises when a mild acoustic pre-stimulus fails to decrease the startle response. Although PPI deficits are a hallmark of schizophrenia [1, 3], they also occur in other psychiatric disorders, such as obsessive compulsive disorder [5, 6], Gilles de la Tourette syndrome [7], Huntington’s disease and post-traumatic stress disorder [8], as well as other neurological disorders such as seizure disorders and nocturnal enuresis [reviewed in ref. 12]. PPI impairments are associated with, and often predictive, of cognitive disruptions and attentional problems. In fact, hallucinations, obsessions, and compulsions are thought to emerge as a result of a deficient gating system that prevents the brain from filtering out irrelevant sensory cues, actions or thoughts [6]. At the present time, the reversal of PPI deficits in animal models and patients with schizophrenia is an efficient tool for antipsychotic drug screening. Despite this, the neuronal pathways and mechanisms underlying the PPI regulatory circuitry are still unresolved. Therefore, identifying the cell types and synaptic mechanisms involved in PPI is crucial to further our understanding of the neuronal underpinning of sensory filtering. Knowledge of the PPI-regulatory circuitry will also have clinical applications, expanding our insights of the pathophysiology of disorders with PPI deficits, towards developing and screening therapeutics for these disorders.

The mammalian startle circuit is simple and consists of the cochlear nuclei which activate giant neurons in the caudal pontine reticular nucleus (PnC) that in turn, directly innervate cervical and spinal motor neurons (MNs) [4, 10, 11] (Fig 1). Previous animal and human studies have shown that inhibition of this pathway by prepulses leads to PPI [1–9]. PPI by acoustic stimuli depends on the activation of midbrain structures including the inferior and the superior colliculi, as well as the pedunculopontine tegmental nucleus (PPTg) [reviewed in ref. 12]. In addition, different cortical and subcortical areas within the corticostriatal-pallido-pontine (CSPP) circuit, regulate PPI. In fact, modulation of the prefrontal cortex, thalamus, hippocampus, basolateral amygdala, nucleus accumbens and dorsal striatum affect PPI [4, 12].

**Figure 1.**
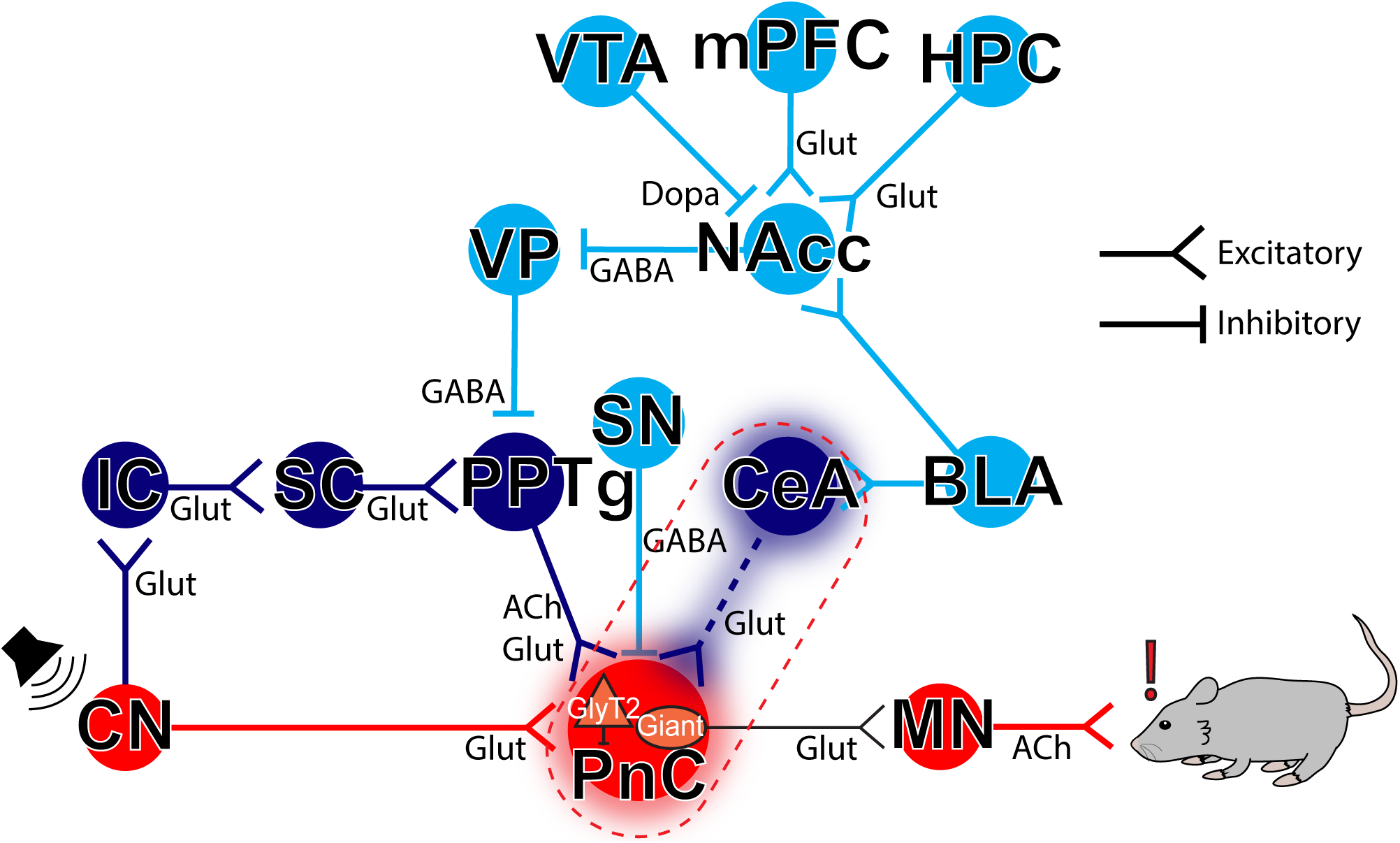
Neuronal circuits contributing to the acoustic startle response and PPI. The mammalian primary acoustic startle pathway (*red pathway*) consists of primary auditory neurons that activate cochlear root and cochlear nuclei (CN), which then relay the auditory information to the giant neurons of the caudal pontine reticular nucleus (PnC) in the brainstem. PnC giant neurons then directly activate cervical and spinal motor neurons (MNs). During PPI (*dark blue pathway*), acoustic prepulses are thought to inhibit startle via the activation of the inferior (IC), superior colliculi (SC) and the pedunculopontine tegmental nucleus (PPTg). The PPI pathway is also under the influence (*light blue pathway*) of midbrain and cortico-limbic structures including the basolateral amygdala (BLA), which activates the nucleus accumbens (NAcc) which in turn, inhibits the ventral pallidum (VP). Together, these PPI structures form the cortico-striato-pallido-pontine (CSPP) network. Here, we propose that CeA-PnC excitatory synapses (*dotted dark blue pathway within the dotted red rectangle*) regulate PPI alongside the CSPP circuit. Glut: Glutamate; Ach: Acetylcholine; Dopa: Dopamine; HPC: hippocampus; mPFC: medial prefrontal cortex; SN: substantia nigra; VTA: ventral tegmental area.

By receiving inputs from various sensory modalities (i.e., trigeminal, vestibular and auditory) and directly activating spinal motor neurons, PnC giant neurons function as key sensorimotor relay neurons within the primary acoustic startle circuit. These PnC giant neurons receive cholinergic inputs from the PPTg, and general lesions of the PPTg were shown to disrupt PPI [13, 14]. Therefore, until recently, there was a consensus that PPI is mediated by cholinergic PPTg neurons inhibiting PnC giant neurons within the startle pathway. However, new optogenetic and chemogenetic rat studies clearly demonstrated that specific activation or inhibition of cholinergic PPTg neurons does not alter PPI [15–17]. Since the majority of neurons in the PPTg are non-cholinergic [18], it is currently suggested that PPI may be a function of GABAergic and/or glutamatergic cells in the PPTg and/or another structure, directly projecting to the PnC. In fact, fish and rodent studies have described the crucial involvement of brainstem glutamatergic, GABAergic and glycinergic inhibitory mechanisms in PPI [19–21].

As it is now clear that PPI does not depend on PPTg cholinergic inputs to the PnC, we wanted to know what structure, other than the PPTg, projects directly onto PnC neurons and modifies PPI. One structure particularly relevant is the central nucleus of the amygdala (CeA), since various neurological disorders showing PPI deficits also show amygdalar abnormalities. In fact, the amygdala is a region that has received considerable attention in studies of the etiology of neuropsychiatric illnesses [22], and impairment of amygdala function disrupts PPI [23–27]. We hypothesized that the CeA-to-PnC connection provides an alternative PPI pathway, independent from the PPTg mechanisms. Our hypothesis is based on *in vivo* and *in vitro* rat studies that corroborate the important role of the CeA in modulating the startle pathway. The CeA receives inputs from the auditory cortex and the central thalamus [28, 29] and sends projections to the PnC [30, 31] making it a potential PPI substrate. More precisely, tract-tracing studies in rats showed that neurons in the rostral part of the medial subdivision of the CeA directly innervate PnC giant neurons at the core of the acoustic startle circuit [31, 32]. However, in contrast to PPI where startle magnitude is *decreased* by a non-startling prepulse, early behavioral studies focused on understanding how startle magnitude is *increased* during conditioned and unconditioned states of fear, as seen both in rats [33, 34] and in humans [35]. These studies, performed in rats, show that the acoustic startle response is enhanced by electrically stimulating the amygdalobasal complex, including the CeA [33] or by injecting NMDA bilaterally in the amygdaloid complex [32]. *In vivo* electrophysiological recordings also revealed that activating the CeA/amygdaloid complex yields excitatory post-synaptic potentials and enhances the acoustic responsiveness of PnC giant neurons in rodents [32, 36].

It is only recently that functional imaging studies and c-Fos expression data in rats have provided strong evidence that CeA neuronal activity is increased during PPI [37]. However, whether and how the CeA-to-PnC excitatory connection, specifically, contributes to PPI has never been tested. Moreover, downstream inhibitory PnC neuronal elements that could help reconcile the role of CeA excitatory projections in a functionally inhibitory pathway, remain to be identified.

Interestingly, glycinergic fibers and interneurons expressing the Glycine transporter type 2 (GlyT2) are closely apposed to the PnC giant neurons in rodents [38, 39]. Although the role and source of activation of these GlyT2^+^ neurons are unclear in the context of PPI, glycine neurotransmission has been shown to inhibit rat PnC giant neurons and contribute to PPI at the level of the startle-initiating Mauthner cells, within the goldfish auditory startle circuit [19, 40].

The present study was undertaken to test the hypothesis that glutamatergic CeA neurons contribute to PPI by sending inputs to the PnC, including GlyT2^+^ neurons in mice. Here we used tract tracing, morphological reconstructions, neurochemical analyses and examined synaptic properties of glutamatergic CeA inputs terminating primarily onto GlyT2^+^ PnC neurons, using transgenic GlyT2-eGFP and GlyT2-Cre mice. We validated our findings *in vivo* by specific optogenetic inhibition and activation of CeA-PnC glutamatergic synapses, as well as optogenetic inhibition of GlyT2^+^ PnC neurons during startle and PPI testing.

## Results

### CeA glutamatergic neurons send inputs to the PnC lateroventral region in mice

As previously reported in rat tracing studies [31, 32], we first confirmed that the retrograde tracer Fluoro-Gold, injected unilaterally within the mouse PnC (Fig. 2A, B), labeled neurons in various brain regions. The cytoarchitectural analysis of the gliotic lesion [41] (Fig. 2, D) showed that Fluoro-Gold was injected within the PnC, delineated by 7th nerve fibers. As expected, Fluoro-Gold labeled neurons in various regions including the caudate putamen, the intra amygdaloid division of the bed nucleus of the stria terminalis, and the lateral hypothalamus (data not shown). Fluoro-Gold also labeled neurons located in the mouse pedunculopontine tegmental area (PPTg) (**Additional file 1: Fig. S1**; N=4) and in the CeA (Fig. 2F; N=4). Labeled CeA cells bodies clustered near the border of the dorsomedial portion of the anterior amygdalar complex, ipsilateral to the PnC injection site.

**Figure 2.**
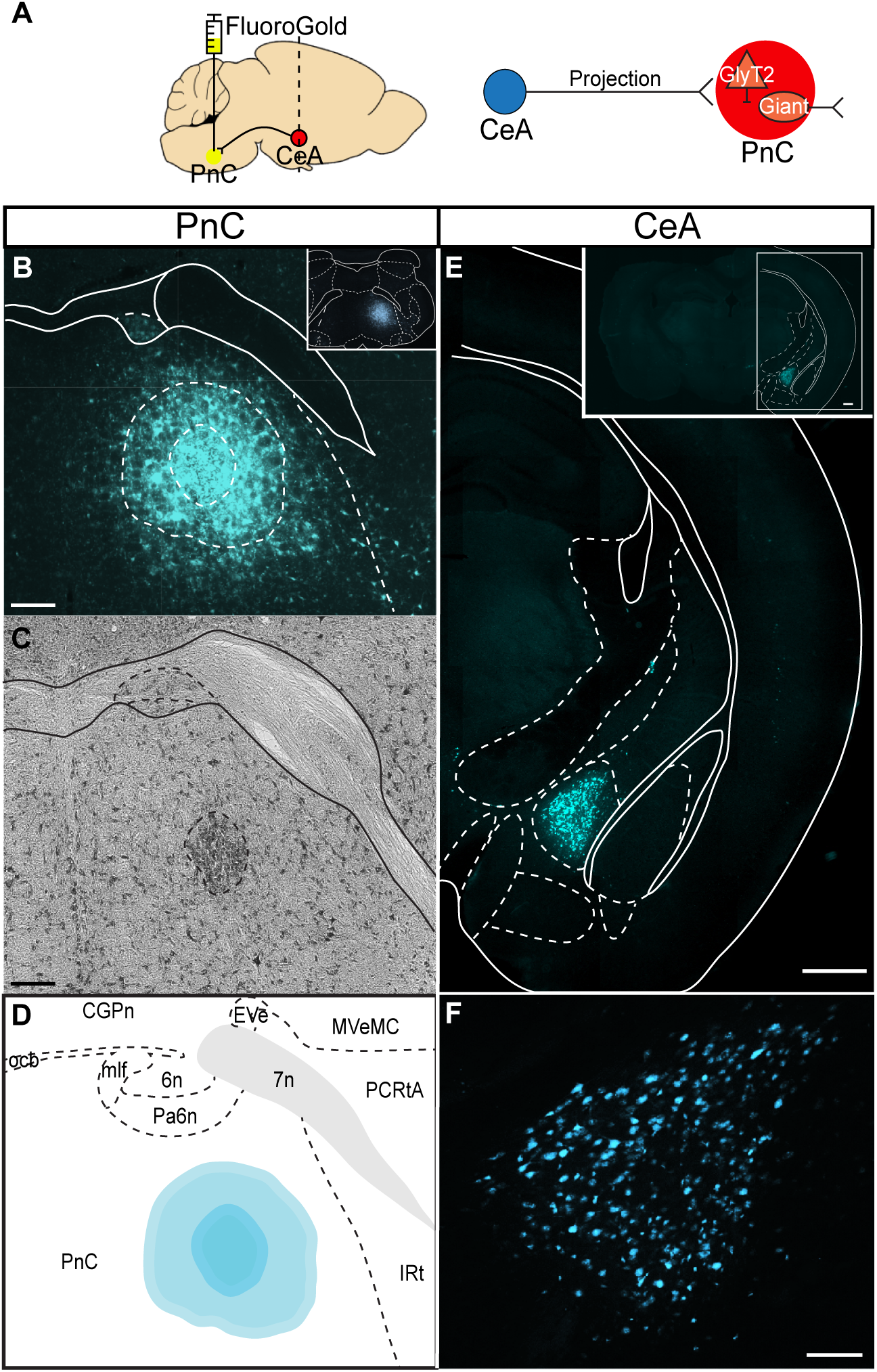
The CeA sends projections to the PnC. (A) *Left*, Sagittal representation of the mouse brain illustrating the Fluoro-Gold injection site in the PnC (yellow circle) and the retrograde labeling site in the CeA (red circle, dotted line). *Right*, Schematic of the hypothesis being tested. (B) Representative coronal PnC slice showing the extent of the Fluoro-Gold injection. The outer dotted circle indicates the fluorescent Fluoro-Gold injection halo. The inner dotted circle represents the center of the gliotic lesion, medial to the 7^th^ cranial nerve and within the cytoarchitectural boundaries of the PnC. *Inset:* Representative image of the injection site in a coronal PnC slice, shown at lower magnification. (C) Representative Nissl-stained PnC section. The darker region surrounded by a dotted circular area indicates the gliotic lesion made by the Fluoro-Gold injection. (D) Representative image showing the Fluoro-Gold injection site in the PnC mapped to the Paxinos and Franklin Mouse Brain Atlas [42]. (E) Representative coronal section showing CeA neurons retrogradely labeled with Fluoro-Gold (cyan). (F) Higher magnification of the CeA neurons shown in (E) and retrogradely labelled with Fluoro-Gold. Representative of N=4 mice. Scale bars: (B-D) 200µm, (E) 500µm, (F) 100µm.

The CeA comprises an array of distinct neuronal populations, including inhibitory neurons that have been classified according to different amygdala markers [43]. The CeA also contains excitatory neurons exhibiting VGLUT2 immunoreactivity, as shown in rats [44]. Since recent evidence suggests that midbrain glutamatergic inputs targeting the pontine startle circuit are important for sensory gating in zebrafish [20, 21], we next focused on how axons from CeA excitatory neurons course within the PnC (Fig. 3**; N=4**). In mice injected with the viral vector AAV-CamKIIα-eYFP into the CeA (Fig. 3A), NeuroTrace™ staining allowed us to confirm that the cell body of eYFP^+^ CeA neurons was efficiently targeted by the viral injection (Fig. 3B-E). Our results show that 11.29% (27 ± 13 somata) of CeA neurons labeled with Neurotrace™ were eYFP^+^ (N=4). Although it is often assumed that CamKIIα is specific to excitatory cells, some GABAergic projection neurons also express CamKIIα [45]. Therefore, we also identified the neurochemical nature of the CamKIIα-eYFP^+^ CeA neurons (Fig. 4A, B) using an *in situ* hybridization assay (RNAscope). We observed that eYFP^+^ CeA neurons express VGLUT2 (Fig. 4C-F, N=3), and, as expected, these neurons were not co-labeled with a GABA antibody (**Additional file 1: Fig. S2**). In fact, within the medial CeA, almost all (83% ± 8%) eYFP^+^ CeA neurons expressed VGLUT2. Finally, we observed that eYFP^+^ CeA axons course predominantly ipsilaterally in PnC sections and are localized in the lateroventral portion of the PnC (Fig. 3F).

**Figure 3.**
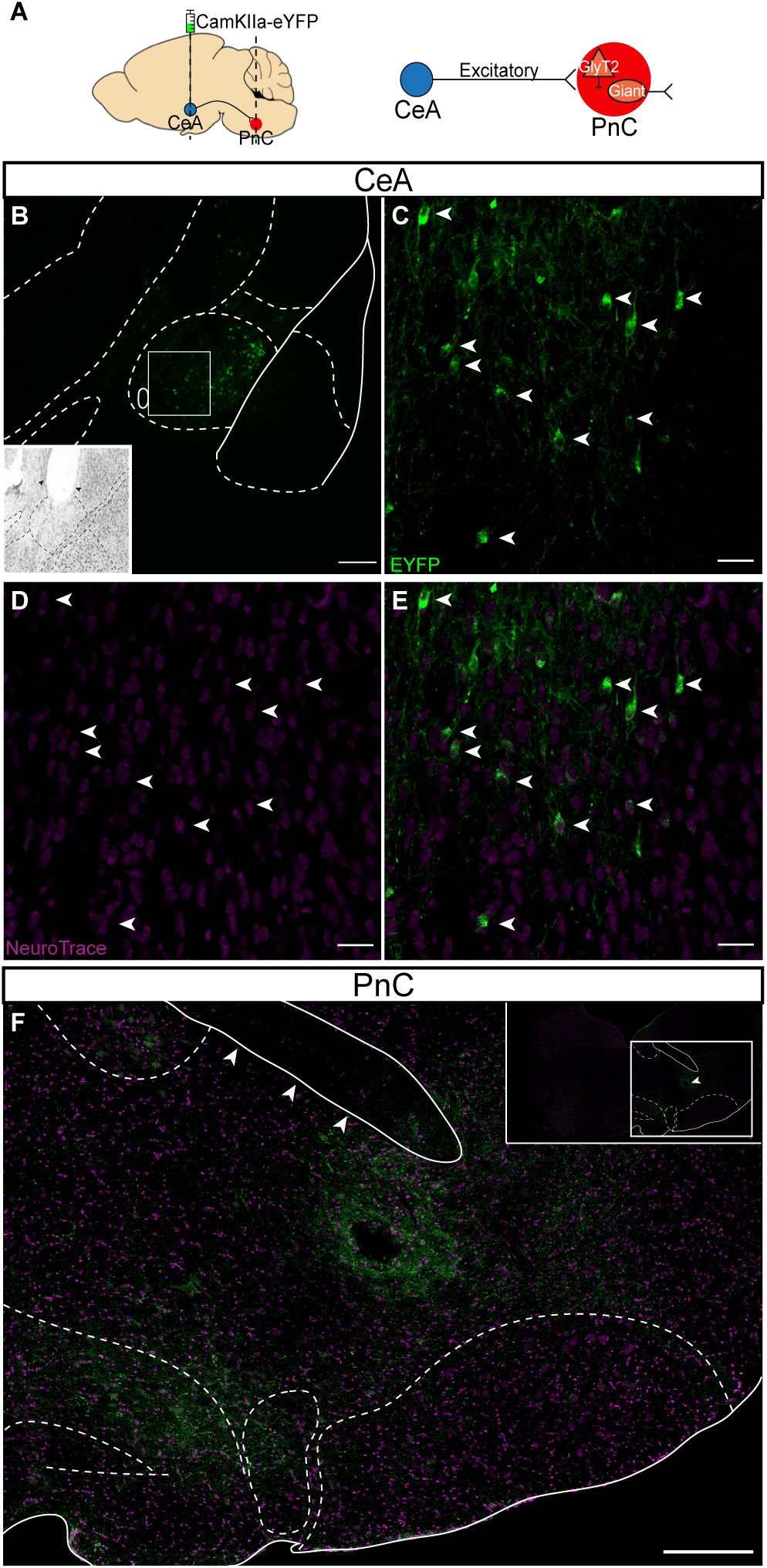
CeA glutamatergic projections course within the lateroventral portion of the PnC. (A) *Left*, Sagittal representation of the mouse brain illustrating the AAV-CamKIIα-eYFP injection site targeting CeA neurons (green circle) and CeA projection fibers terminating at the level of the PnC (red circle). The dotted line illustrates the PnC level at which coronal cut sections were obtained to visualize CamKIIα-eYFP^+^ axons originating from the CeA. *Right*, Schematic of the hypothesis being tested. (B) Representative CeA coronal sections showing eYFP^+^ fluorescence (green) and NeuroTrace™ staining (magenta). The white rectangle shows the area imaged in panels C-E. *Inset*, Nissl stain image of injection site in the CeA. Arrowheads represent the cannula tract. (C) White arrows indicate CeA cells positive for CamKIIα-eYFP (green). (D) NeuroTrace™ stain (magenta) labels CeA cells bodies (white arrows) (E) White arrows indicate CeA cells positive for CamKIIα-eYFP and NeuroTrace™. (F) Representative image of CamKIIα-eYFP^+^ CeA fibers (green) coursing within a PnC coronal section stained with NeuroTrace™ (magenta). *Inset*, Representative image of a PnC slice showing the tract (arrow) of the implanted optic fiber for PPI *in vivo* experiments. Representative of N=4 mice. Scale bars: (C, E) 400µm, (D-F) 50µm.

**Figure 4.**
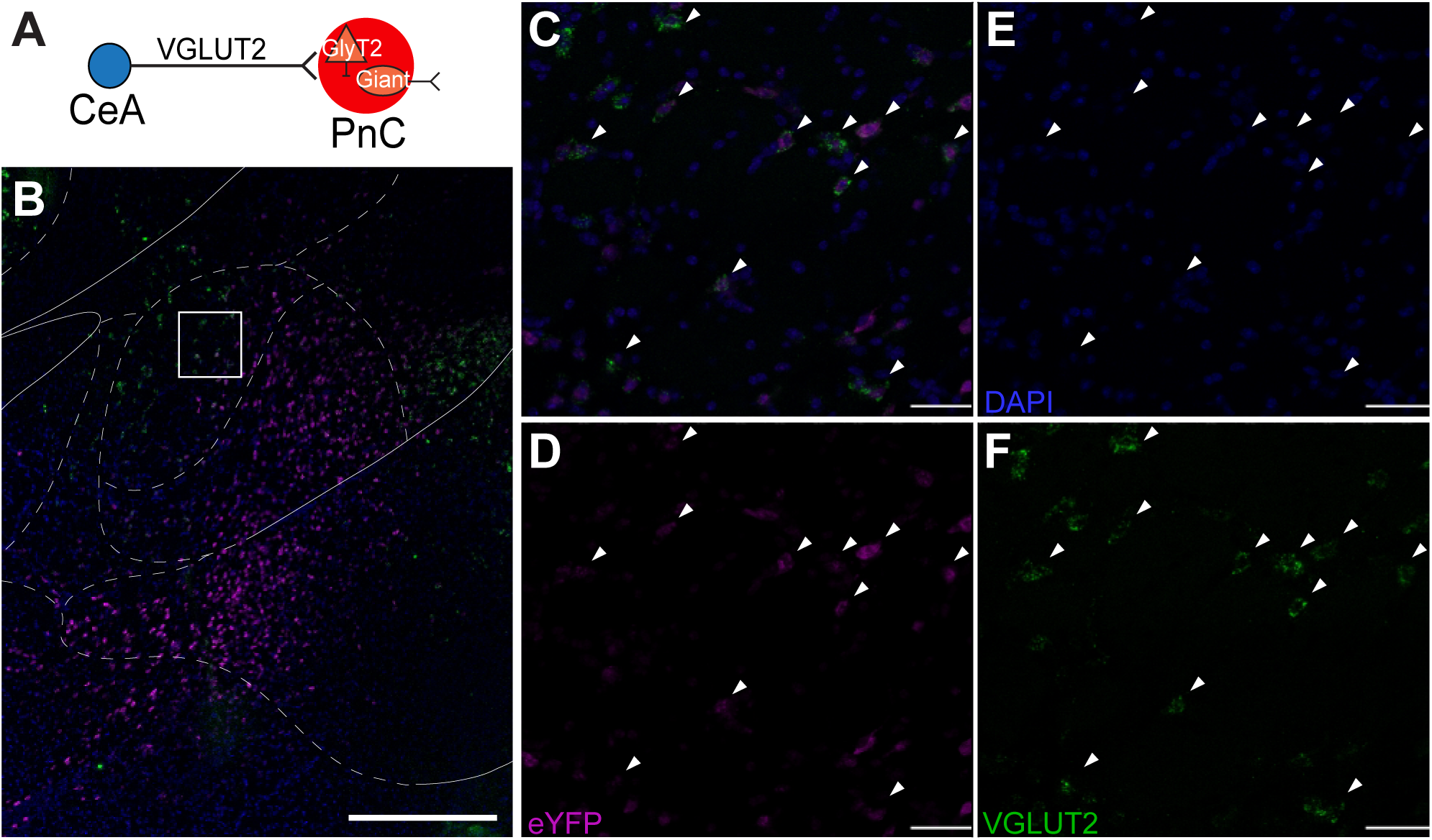
CeA neurons targeted with the AAV-CamKIIα-eYFP viral injection are VGLUT2^+^. (A) Schematic of the hypothesis being tested. (B) Representative image of a CeA coronal section at low magnification, hybridized with eYFP (magenta) and VGLUT2 (green) probes. White rectangle shows area imaged in panels C-F. (C-F) Arrowheads indicate CamKIIα-eYFP^+^ CeA neurons co-expressing VGLUT2 mRNA. Representative of N=3 mice. Scale bars: (B) 500 µm, (C-F) 25 µm.

### Optogenetic inhibition of CeA-PnC excitatory synapses during acoustic prepulses decreases PPI

Following our histological analyses, we assessed the potential role of the CeA-PnC excitatory connection during PPI, *in vivo*. We anticipated that silencing amygdala neurons that regulate PPI would not affect baseline startle in the absence of a prepulse. To silence CeA-PnC excitatory synapses, we used WT mice injected with the optogenetic viral vector rAAVDJ/CamKIIa-eArch3.0-eYFP to transduce CeA excitatory cells with Archaerhodopsin-3 (Arch3.0). We tested non-injected mice (control) and mice injected with the control rAAVDJ/CamKIIa-eYFP for comparison. Following the unilateral intracranial injection, an optic fiber was implanted at the level of the PnC, to optically inhibit CeA fibers/terminals expressing Arch3.0 (Fig. 5A**, left**).

**Figure 5.**
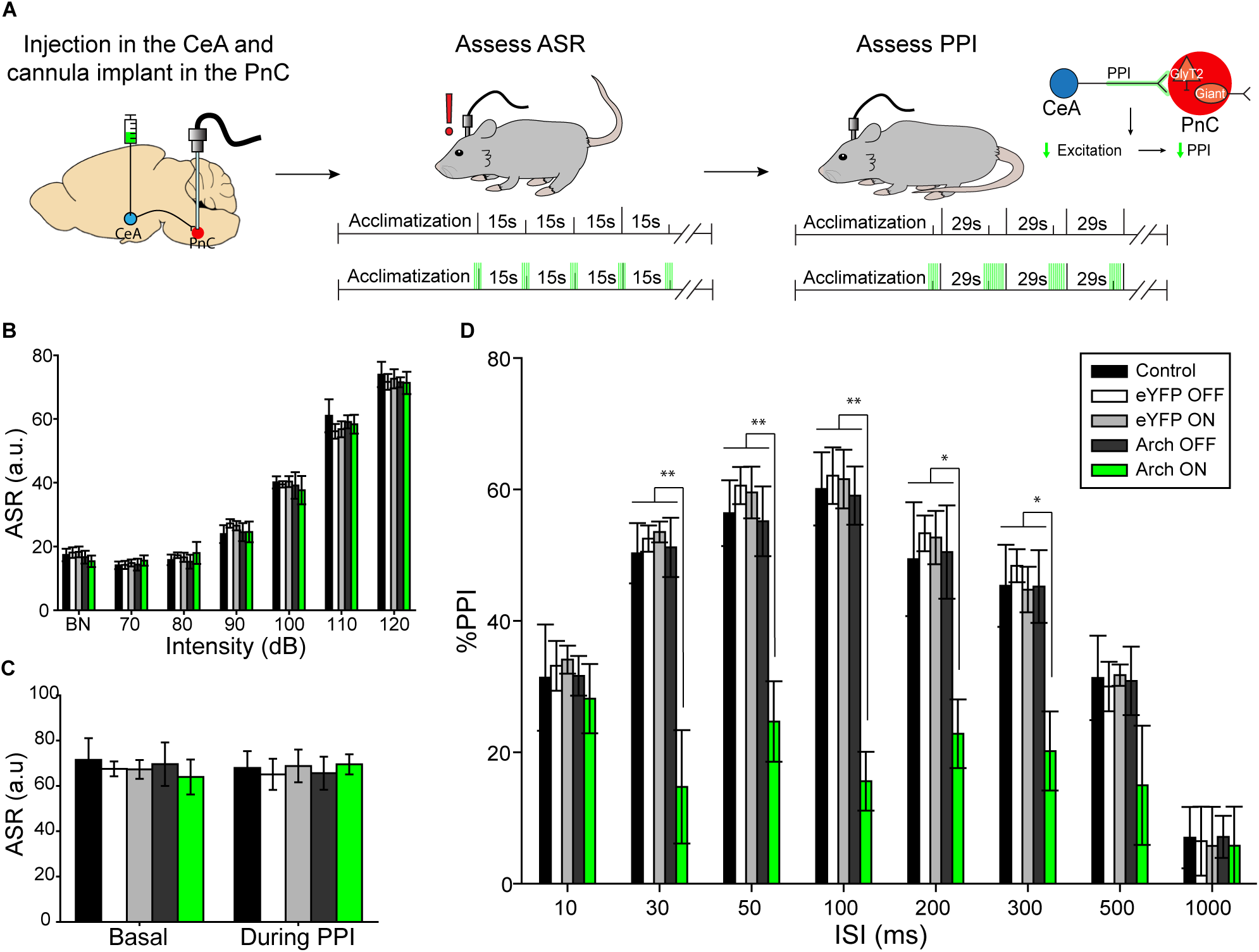
Silencing CeA-PnC excitatory projections during acoustic prepulses decreases PPI. (A) Schematic of acoustic startle reflex and PPI protocols performed using non injected WT control mice, mice injected with eYFP only (light ON or OFF) and mice injected with Archaerhodopsin (Arch3.0; light ON or OFF). The rightmost schematic represents the hypothesis being tested. (B) Graph showing no significant effect of green light paired with 70dB-120dB acoustic startling pulses on basal startle amplitude among animal groups [mouse group: (F_(1,11)_=1.417, p=0.268); light: (F_(1)_=0.00155, p=0.969); sound intensity*light interaction: (F_(1,6)_=0.206, p=0.974)]. (C) Graph showing no significant main effect of light during 120dB pulses presented before (basal) vs. randomly during the PPI task, on mean baseline startle amplitude among animal groups (F_(1)_=3.124, p=0.105). (D) Graph showing that green light paired with acoustic prepulses significantly decreased PPI only in mice injected with Arch3.0, at ISIs between 30 and 300ms. We found a significant effect of ISI (F_(1,7)_=24.863, p<0.001), light: (F_(1)_=10.201, p=0.009), and light*ISI interaction: (F_(1,7)_=4.057, p<0.001) on PPI (Two-way RM ANOVA). N=8 mice per group. Data are represented as mean ± SEM. *p<0.05, **p<0.01.

To test our hypothesis that silencing CeA-PnC excitatory synapses does not affect baseline startle elicited by a pulse alone stimulation (“pulse”), we photo-inhibited CeA-PnC excitatory synapses prior to and concurrent with the acoustic stimulation at increasing sound levels, and then, we measured the startle response as a readout (Fig. 5A**, middle**). In all mice, sound intensities of and beyond 90 dB led to a measurable acoustic startle reflex, characterized by a whole-body flexor muscle contraction [2, 10, 11]. We found no differences when we compared the startle amplitude obtained with and without photo-inhibition of CeA-PnC excitatory synapses from eArch3.0-expressing animals to animals injected with the control virus and non-injected controls (Fig. 5B). These results, which were replicated in mice injected with Halorhodopsin (pAAV DJ-CamKIIa-NpHR3.0-eYFP; **Additional file 1: Fig. S3A-B**), confirm that inhibiting CeA-PnC excitatory synapses prior to and during a startle stimulation does not affect baseline startle. Our observations are consistent with previous rat studies demonstrating that lesions of the CeA do not block the acoustic startle response itself [46].

We then tested whether silencing CeA-PnC excitatory synapses affects PPI trials. To do so, the photo-inhibition started with prepulse onset, and lasted until the end of the inter-stimulus intervals (ISIs) between prepulse and pulse (Fig. 5A**, right**) [15–17, 20]. Although photo-inhibition had no impact on pulse-alone stimulations interspersed with PPI trials (Fig. 5C), photo-inhibition of CeA-PnC excitatory connection during the prepulse significantly decreased PPI by 25-43% at ISI between 30ms-300ms in eArch3.0-expressing animals, but not in the control animals (Fig. 5D). Similarly, in mice injected with Halorhodopsin, photo-inhibition significantly decreased PPI by 16-29% when the prepulse was presented 50-300ms before the startling pulse (**Additional file 1: Fig. S3C**), without altering startle magnitude in pulse-alone trials (**Additional file 1: Fig. S3A-B**). Since silencing the CeA-PnC excitatory connection during the prepulse and the subsequent ISI led to a decrease in PPI, these results support our hypothesis that CeA excitatory neurons regulate part of the behavioral PPI.

Next, we tested whether photo-activation of CeA-PnC excitatory synapses prior to a pulse-alone stimulation could mimic the effect of an acoustic prepulse and lead to PPI (Fig. 6A). To test this, control mice injected with eYFP only were used to test for a possible light-induced heat effect. These mice were compared to WT mice injected with the optogenetic AAV vector rAAVDJ-CamKIIα-ChR2-eYFP in the CeA. In all mice, CeA-PnC excitatory synapses were photo-activated shortly prior to a startling pulse, at intervals used for PPI. As shown in Figure 6B, activating this connection with blue light pulses at 5Hz or 20Hz produced a PPI effect across different ISI (between photo-activation and acoustic pulses) only in WT mice injected with ChR2. Interestingly, the 20Hz photo-activation tended to be more efficient than the 5Hz photo-stimulation train (ISI=50ms, q=4.376, p=0.007), yielding PPI values 18-41% of PPI elicited by the acoustic prepulse. Overall, the light-induced PPI effect was smaller than PPI induced by an acoustic prepulse, which resulted in a 35-60% PPI. Importantly, our results confirm that CeA excitatory neurons sending inputs to the PnC contribute to PPI, during the interval between the prepulse and the pulse.

**Figure 6.**
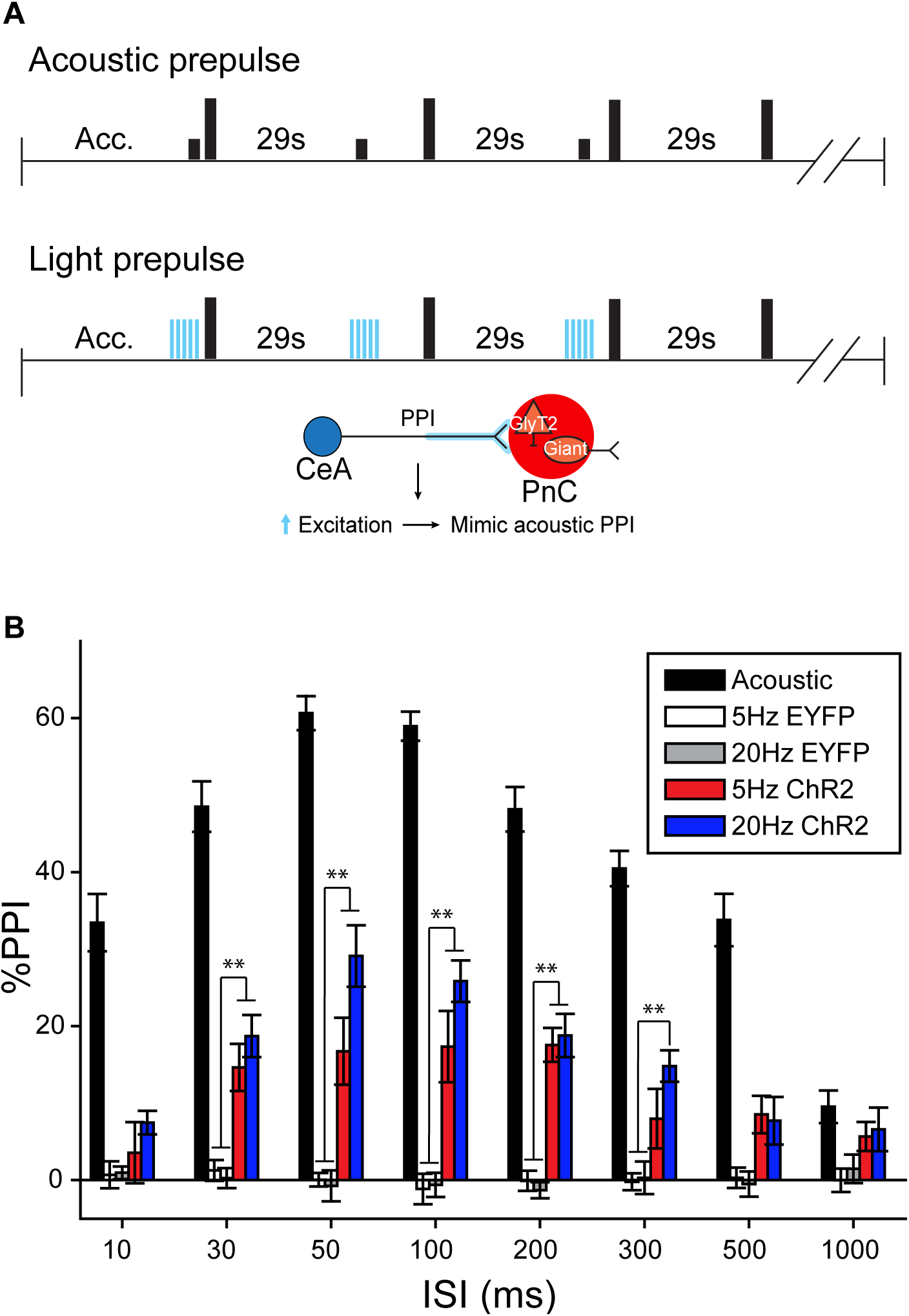
Photo-stimulation of CeA-PnC excitatory synapses induces PPI. (A) Representation of the PPI protocols performed using acoustic prepulses (*top*) or blue light prepulses (*middle*). The *bottom* schematic represents the hypothesis being tested. (B) PPI was assessed using acoustic prepulses (black bars) or brief optogenetic stimulation of CeA-PnC glutamatergic synapses at 5Hz (red bars) and 20Hz (dark blue bars) in WT mice injected with ChR2-eYFP and control mice injected with eYFP only (white and grey bars) used to test for a possible light-induced heat effect. Graph showing that optogenetic stimulation elicited PPI values (i.e., 18-41% of acoustic prepulse) lower than PPI values obtained using acoustic prepulses, at ISI between 10-500ms. N=8 mice per group; ANOVA (F_(1,14)_=6.152, p<0.001).

Finally, we tested whether photo-stimulation of CeA-PnC excitatory synapses (at 20 Hz) enhances the effect of the acoustic prepulse and potentiates PPI. Therefore, we paired the photo-stimulation with pulse-alone stimulations or with acoustic prepulses during PPI trials, using mice injected with AAV-CamKIIα-ChR2-eYFP (**Additional file 1: Fig. S4;** N=6). Photo-stimulation did not alter startle responses (**Additional file 1: Fig. S4A;** p>0.05) or PPI (**Additional file 1: Fig. S4B**; p>0.05). Based on our results, we conclude that CeA-PnC excitatory synapses are maximally activated in response to acoustic prepulses *in vivo*, and that additional photo-stimulation of CeA-PnC excitatory inputs during acoustic prepulses does not further enhance PPI.

### CeA glutamatergic neurons send inputs to PnC inhibitory neurons

We next aimed to reconcile the glutamatergic nature of CeA-PnC inputs and their role in a functionally inhibitory pathway. Since glycinergic neurons are in the PnC and in close proximity of PnC giant neurons, we hypothesized that CeA excitatory inputs activate glycinergic PnC neurons, contributing to the inhibitory component of the PPI phenomenon. To test our hypothesis, we injected the AAV viral vector AAVDJ-CamKIIα-mCherry into the CeA of transgenic mice expressing eGFP under the control of the glycine transporter 2 (GlyT2) promoter [39]. Orthogonal imaging and morphological reconstruction analyses revealed close putative synaptic appositions between CamKIIα-mCherry^+^ CeA projections and GlyT2^+^ PnC neurons (Fig. 7A-C; **N=6**). In addition, PSD-95, a post-synaptic protein at excitatory synapses, co-localized with these appositions confirming that CamKIIα-mCherry^+^ CeA fibers form excitatory synapses with GlyT2^+^ PnC neurons (Fig. 7C).

**Figure 7.**
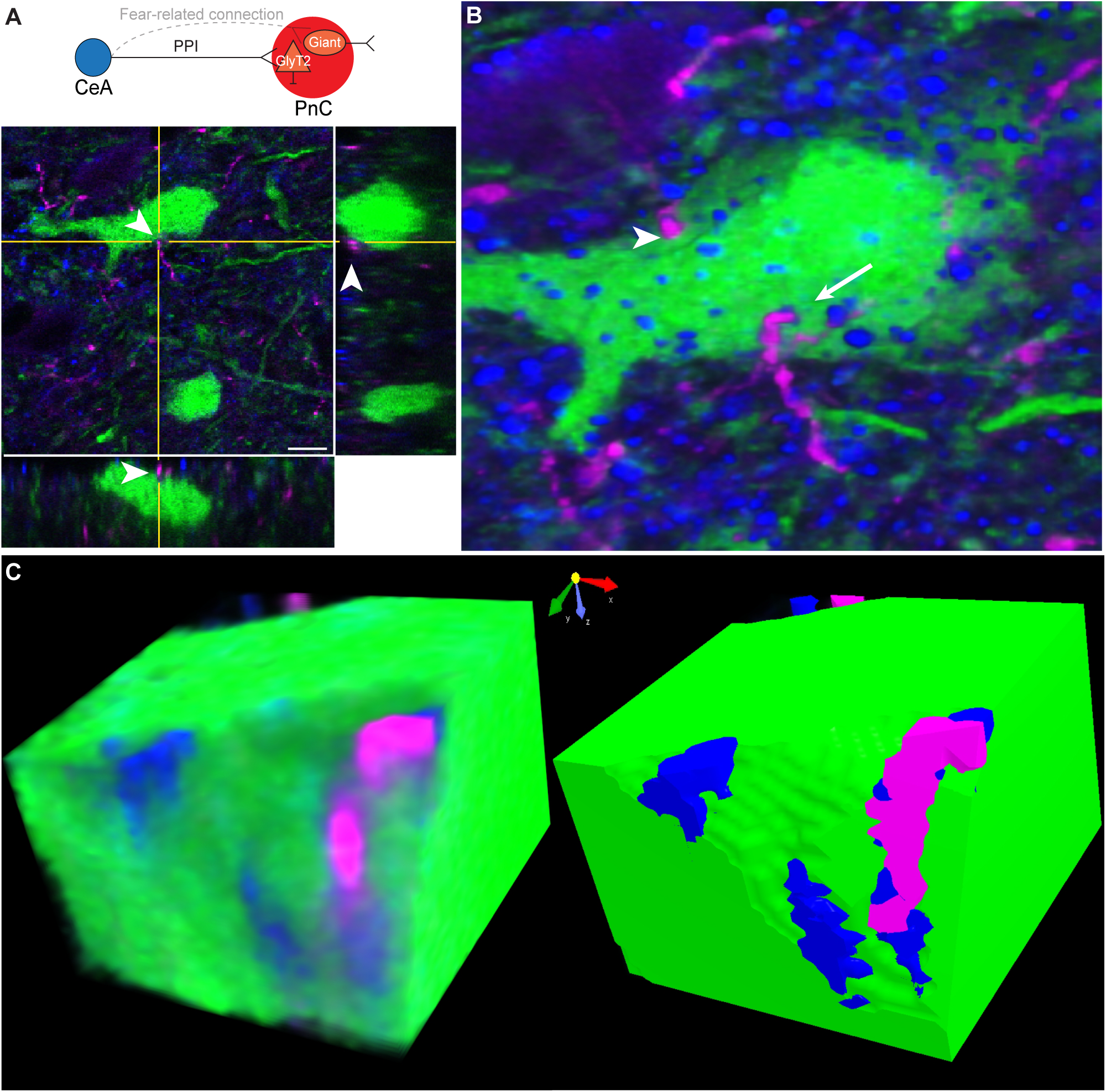
CamKIIα^+^ CeA fibers closely apposed to GlyT2^+^ PnC neurons. (A) *Top*, Schematic of the hypothesis being tested. *Bottom*, Orthogonal view of a close apposition between CamKIIα-mCherry^+^ CeA excitatory fibers (magenta) and the soma of a GlyT2^+^ PnC neuron (green) indicated by the arrowhead in all three views. (B) Three-dimensional reconstruction of putative synaptic contacts between CamKIIα-mCherry^+^ CeA fibers (magenta) and GlyT2-eGFP^+^ neurons expressing PSD-95 (blue; arrow). Some putative synaptic appositions did not show PSD-95 staining (arrowheads). (C) Volume rendering and angular sectioning of the PSD-95^+^ putative synaptic contact shown in B. Scale bar in (A) is 50µm.

Next, we recorded basic electrical properties of CamKIIα-eYFP^+^ CeA neurons expressing ChR2 while confirming their sensitivity to blue light photo-stimulation. For this, *in vitro* patch clamp recordings were performed in acute CeA slices of mice expressing the Cre-recombinase enzyme in GlyT2^+^ neurons (i.e., GlyT2-Cre mice). These GlyT2-Cre mice were previously injected with the AAVDJ-CamKIIα-ChR2-eYFP viral vector in the CeA and a Cre-dependent tdTomato viral vector in the PnC, to transduce CeA excitatory cells with ChR2 and GlyT2^+^ PnC neurons with tdTomato, respectively (Fig. 8A **and Additional file 1: Fig. S5A**). Spontaneous excitatory post-synaptic currents (sEPSC) with a mean amplitude of 13.26 ± 0.1 pA were recorded in CamKIIα-eYFP^+^ CeA neurons held at −70mV, and current injections elicited action potentials firing at a maximum rate of 14.6 ± 5.4 Hz (**Additional file 1: Fig. S5C**). More importantly, photo-stimulation induced large current responses (821.15 ± 20.3pA maximum amplitude), indicating that our stimulation protocol successfully activated CamKIIα-eYFP^+^ CeA neurons expressing ChR2 (**Additional file 1: Fig. S5D**).

**Fig 8.**
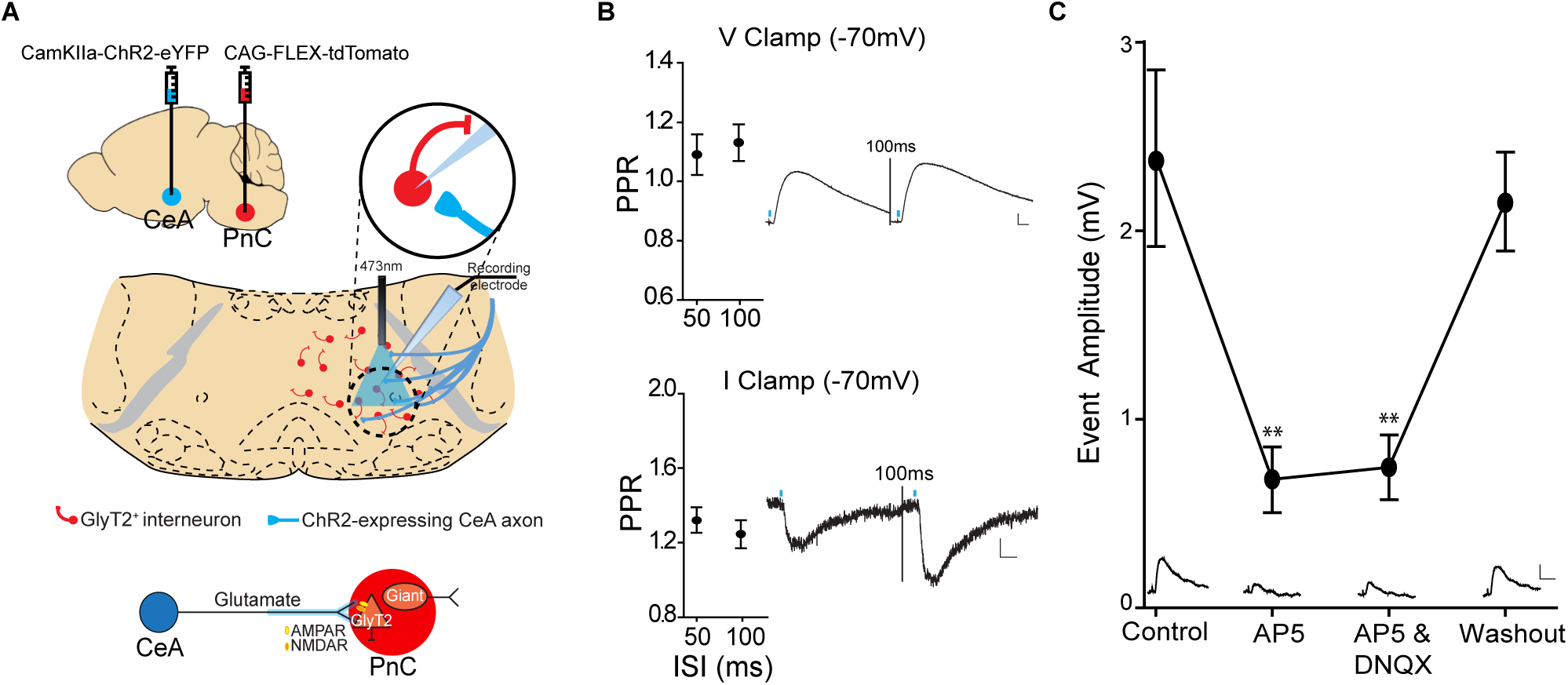
CeA glutamatergic inputs activate GlyT2^+^ PnC neurons via AMPA and NMDA receptors. (A) *Top*, Injection of AAVDJ-CamKIIα-ChR2-eYFP in the CeA and injection of Cre-dependent AAVDJ-tdTomato in the PnC of GlyT2-Cre mice, followed by *in vitro* patch clamp recordings. *Bottom*, Schematic of the hypothesis being tested. (B) Paired-pulse ratios of the light-evoked EPSPs (*Top*) and EPSCs *(Bottom)* at 50 and 100ms interstimulus intervals (ISIs). *Insets*: Representative traces. (C) Graph showing the amplitude of the light-evoked EPSPs recorded in GlyT2^+^ PnC neurons in control, during the sequential bath-application of AP5 and DNQX and following washout. *Insets*: Sample traces. Scale: 10mV/15ms. Representative of N=10 mice, n=38 neurons. Data are represented as mean ± SEM. *P>0.05, **P>0.01. Scale bars: (B) Voltage traces: 2mV/10ms; Current traces: 5pA/5ms

We then used acute PnC slices obtained from the same mice to determine whether photo-stimulation of CeA excitatory fibers could elicit excitatory synaptic responses in GlyT2^+^ PnC neurons (Fig. 8A). Photo-stimulation (i.e., blue light pulses) evoked excitatory post-synaptic potentials (EPSPs) and currents (EPSCs) in td-Tomato-expressing GlyT2^+^ neurons held at −70mV. These excitatory responses showed facilitation at ISI of 50 and 100ms (Fig. 8B, EPSP PPR: 50ms=1.09, 100ms=1.13; EPSC PPR: 50ms=1.318, 100ms=1.243). Then, we applied glutamate receptor antagonists to functionally identify the post-synaptic receptors underlying the EPSPs (2.39 ± 0.47mV) elicited in GlyT2^+^ PnC neurons, in response to the CeA fibers photo-stimulation. AP5 (50μM) eliminated the NMDAR-dependent component of the EPSP (AP5: 0.68 ±0.17mV; 1-way RM ANOVA, F=9.463, p<0.01) and DNQX (20μM) blocked the (AMPAR)-dependent component (Fig. 8C; DNQX: 0.75 ± 0.17mV; 1-way RM ANOVA, F=6.009, p<0.01) which recovered by washing out the drugs (2.15 ± 0.75mV; Fig. 8C).

Next, to determine if the electrical properties of GlyT2^+^ PnC neurons targeted by CeA excitatory inputs differ from neighboring untargeted GlyT2^+^ PnC cells (Fig. 9A), we compared the intrinsic and spontaneous synaptic properties of tdTomato-expressing GlyT2^+^ neurons (**Additional file 1: Fig. S6 and Table S1**) responsive to light vs. tdTomato-expressing GlyT2^+^ neurons unresponsive to light, at −70 mV. Their anatomical location was confirmed post-recording using GlyT2 and Biocytin co-immunostaining, followed by a 3D reconstruction (Fig. 9B-C). As expected from our tract-tracing results, GlyT2^+^ cells medial to 7n (n=6) and cells located lateroventrally were responsive to blue light (n=12; Fig. 9B-C) whereas cells located along the midline (n=16) or contralateral to the injection site (n=4) did not respond to light (not shown). The EPSPs and EPSCs recorded at −70 mV were abolished at 0 mV (i.e., the EPSP reversal potential). None of the light-responsive cells showed IPSP or IPSCs at 0 mV, confirming that no inhibitory inputs were activated by blue light (n=12; Fig. 9D). While both cell types displayed similar passive and active membrane properties (**Additional file 1: Fig. S7**) (F=1.119, p=0.327; **Additional file 1: Fig. S6C**) and received spontaneous excitatory and inhibitory inputs, the amplitude of sEPSCs (n=18, t=2.538, p=0.011) and sIPSCs (t=2.434, p=0.025) of GlyT2^+^ responsive cells was greater compared to that of GlyT2^+^ unresponsive cells (n=20; **Additional file 1: Fig. S6-B**).

**Figure 9.**
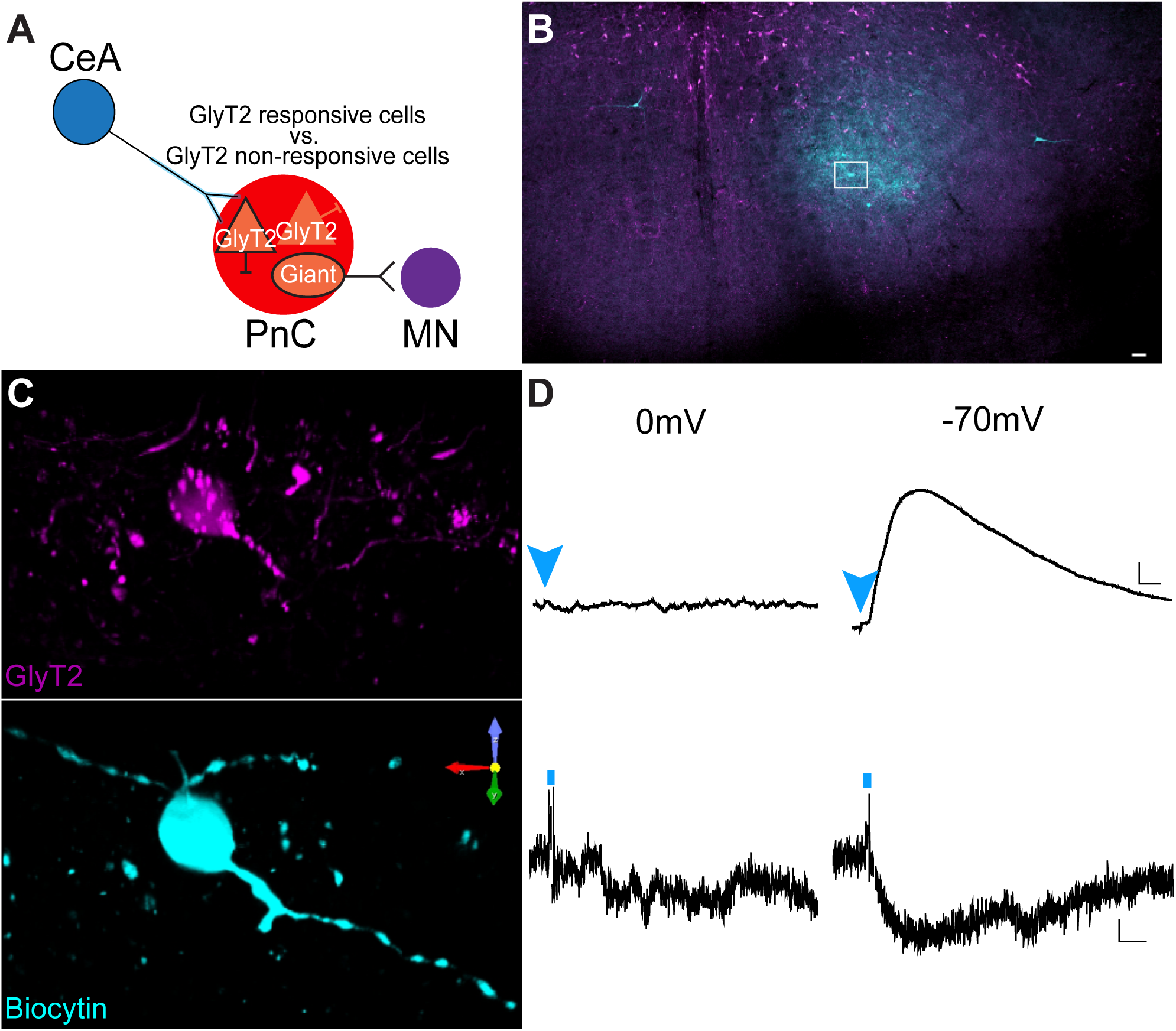
Electrophysiological properties of GlyT2^+^ PnC neurons. (A) Schematic of the hypothesis being tested. (B)Representative PnC slice showing eGFP^+^ fluorescence (magenta) and Biocytin staining (cyan). (C) Higher magnification of the box area in (A), showing representative morphological reconstructions of recorded GlyT2^+^ cell bodies filled with biocytin. (D) Representative light-evoked voltage (top two traces) and current traces (bottom two traces) recorded at 0mV (left) and −70mV (right) of a GlyT2^+^ neuron responsive to blue light. Blue arrowheads and lines indicate blue-light photo-stimulation. Scale bars: (B) 250 µm. (C) 10 µm. (D) Voltage traces: 1mV/10ms; Current traces: 5pA/1ms.

Results of these electrophysiological experiments confirm our anatomical data showing that CeA glutamatergic projections activate a subset of GlyT2^+^ PnC neurons located latero-ventrally, via AMPA and NMDA receptors. Spontaneous IPSCs were recorded in GlyT2^+^ PnC neurons held at 0mV (EPSP reversal potential), confirming that these neurons receive inhibitory inputs. However, since blue light photo-stimulation failed to evoke IPSPs in GlyT2^+^ PnC neurons, these results confirm that CamKIIα^+^ CeA neurons do not send inhibitory projections to GlyT2^+^ PnC neurons.

### Optogenetic inhibition of GlyT2^+^ PnC neurons during acoustic prepulses decreases PPI

Finally, to test whether GlyT2^+^ PnC neurons (likely activated by CeA inputs) contribute to PPI (Fig. 10A), we studied the behavioral contribution of GlyT2^+^ PnC neurons, using GlyT2-Cre mice (N = 8 mice) injected with the Cre-dependent optogenetic viral vector rAAVDJ/Ef1α-DIO-eArch3.0-eYFP to transduce GlyT2^+^ PnC neurons with Archaerhodopsin-3 (Arch3.0). We optogenetically inhibited these neurons during PPI through unilateral optic fibers, chronically implanted in the PnC. Photo-inhibition of GlyT2^+^ PnC neurons using 1ms pulses of green light stimulation presented at 5Hz had no impact on the acoustic startle response (Fig. 10B) or pulse-alone stimulations interspersed with PPI trials (Fig. 10C). However, photo-inhibition of GlyT2^+^ PnC neurons during the prepulse significantly decreased PPI by 37%-40% at ISI between 30ms-100ms (Fig. 10D).

**Figure 10.**
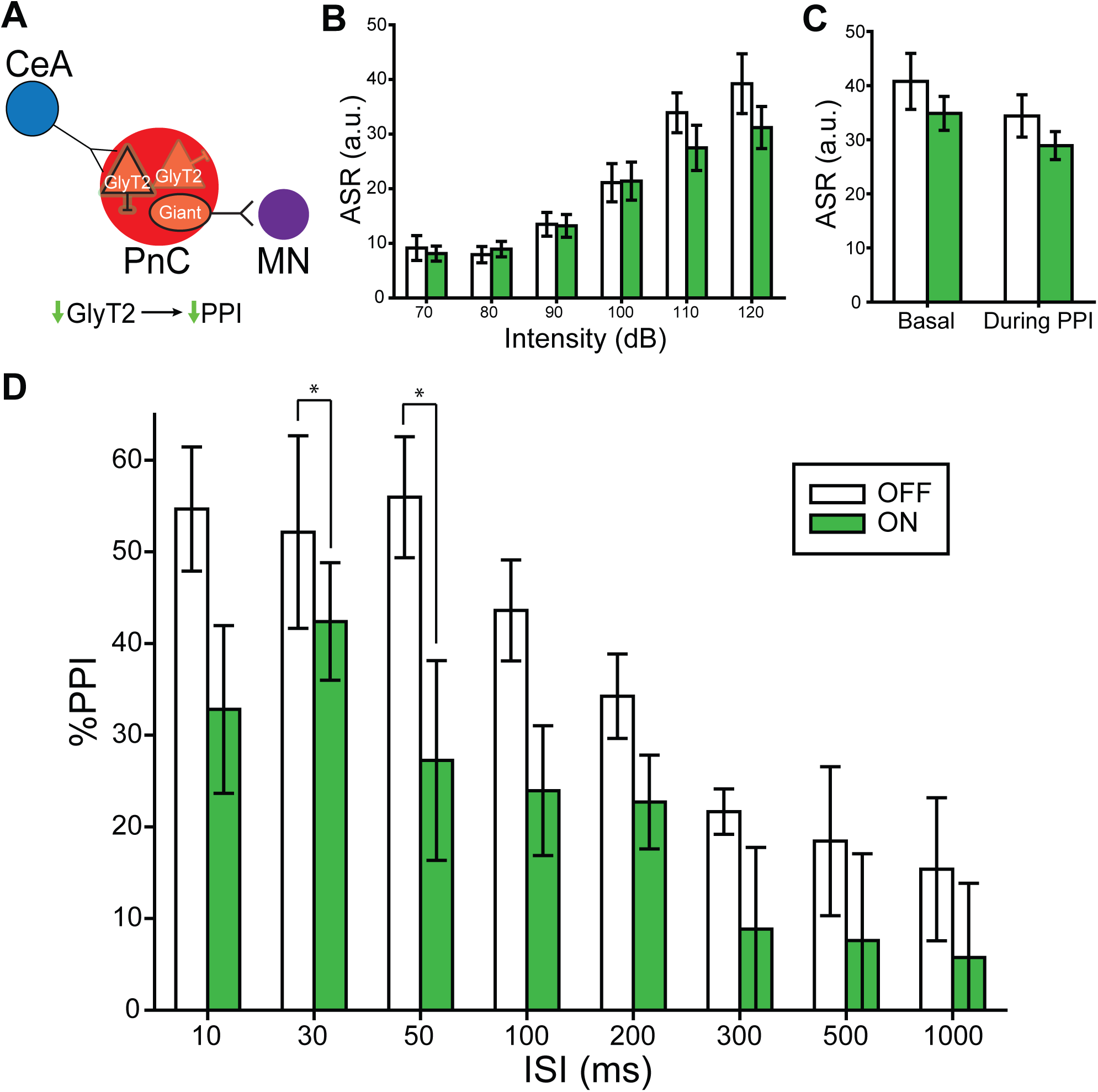
Silencing GlyT2^+^ PnC neurons during acoustic prepulses decreases PPI. (A) Schematic of the hypothesis being tested in GlyT2-Cre mice injected with a Cre-dependent AAV encoding Archaerhodopsin-eYFP (Arch3.0-eYFP). (B) Graph showing no significant effect of green light paired with 70dB-120dB acoustic startling pulses on basal startle amplitude [light: (F_(1,7)_=1.407, p=0.274); intensity*light interaction:(F_(3,21)_ =1.747, p=0.188)]. (C) Graph showing no significant main effect of light during 120dB pulses presented before (basal) vs. randomly during the PPI task, on mean baseline startle amplitude (F_(1)_=3.124, p=0.105). (D) Graph showing that green light paired with acoustic prepulses significantly decreased PPI in mice injected with Arch3.0, at ISIs between 30 and 100ms. We found a significant effect of ISI (F_(6,42)_=8.957, p<0.001), and light (F_(1,7)_=8.216, p=0.024).(Two-way RM ANOVA). N=8 mice per group. Data are represented as mean ± SEM. *p<0.05, **p<0.01.

## Discussion

Overall, our results confirm that in mice, the PnC receives CeA glutamatergic projections which course predominantly ipsilaterally, within the ventrolateral portion of the PnC. Since photo-activating these excitatory projections at the level of the PnC induced a partial PPI and photo-silencing this connection produced an ISI-dependent reduction in PPI, this suggests that CeA glutamatergic inputs contribute to PPI. We also found that the GlyT2^+^ PnC neurons responsive to the photo-stimulation of CeA glutamatergic inputs display AMPA and NMDA receptor-dependent excitatory responses. Finally, our results show that silencing GlyT2^+^ PnC neurons decreases PPI. Together, the influence of CeA glutamatergic neurons in PPI and their direct input to GlyT2^+^ neurons, located at the core of the PnC startle circuit, strongly argue that CeA glutamatergic neurons and GlyT2^+^ neurons are an intrinsic part of the neuronal mechanism underlying the prepulse inhibition of the startle response.

### Anatomical studies

Our data confirm that in mice, there is a direct pathway originating from the CeA onto the pontine reticular formation, as previously described in rats, cats, guinea pigs and monkeys [44, 47–51]. Our retrograde tracing experiments revealed that afferent projections to the PnC originate in various brain regions including the PPTg and the CeA (Fig. 2 **and Additional file 1: Fig. S1**). Recent evidence in rats and fish suggest that glutamatergic (and likely also, GABAergic cells), but *not* cholinergic neurons, in the PPTg are essential for inhibiting the startle response, during PPI [15-17, 20, 21]. Here, we focused on the glutamatergic projections from the CeA because this region was found to be both relevant for the modulation of the acoustic startle response [32–34] and relevant to diseases associated with sensory gating deficits [37]. In rats, while PnC-projecting CeA neurons were shown to be able to enhance startle reactivity of PnC giant neurons, immunohistochemical assays revealed their non-GABAergic nature [52]. In contrast to the well described CeA inhibitory neuronal populations [43], very few immunohistochemical studies have investigated the population of CeA neurons expressing the glutamatergic neuronal marker VGLUT2^+^, which directly project to pontine neurons. Fung et al. (2011) reported that 24% of all retrogradely-labeled CeA neurons that project directly to the oral pontine reticular nuclei (PnO; rostral to the PnC) are VGLUT2^+^ and are located in the lateral and capsular subdivisions of the CeA. Although the studies of Fung et al (2011) were conducted in guinea pigs, our anatomical data obtained in mice are similar: that is, our CamKIIα-driven tract-tracing analysis and *in situ* hybridization results (Figs. 3 and 4) also confirm that 80% of the CamKII^+^ CeA neurons virally targeted, are glutamatergic and project to the PnC. Interestingly, we demonstrate that amygdalar cell bodies sending projections to the PnC are confined to the medial CeA; and no cell bodies in the lateral and capsular CeA were detected.

Our anterograde tracing experiments and histological analyses also show that the descending CeA glutamatergic fibers course into the ventro-lateral part of the PnC, adjacent to the 7th nerve fibers and the olivary complex, in mice (Fig. 3F). Our findings are in accordance with the results of previous anatomical studies in rats and cats showing that descending CeA projecting neurons innervate, directly, with ipsilateral predominance, neurons in the PnC [47–51]. The lateroventral PnC is innervated by cochlear nuclei fibers conveying acoustic startle input to PnC neurons, as lesions in the lateroventral PnC greatly attenuate the startle reflex [10, 11, 53]. Altogether these findings confirm that the CeA projects to a region in the PnC essential for acoustic startle processing.

Interestingly, previous analysis had shown that CeA terminal fibers were only occasionally seen close to the *dendrites* of PnC giant neurons responsible for the startle response [25], in contrast to projections from the cochlear nucleus, which terminate on the *somata* and proximal dendrites of these PnC giant neurons [54; Zaman et al., 2017]. Rather, such CeA fibers mainly terminated close to the *somata* of small and medium-sized neurons of the PnC, whose chemical phenotype was not identified [32]. The small diameter (10-20um) GlyT2^+^ PnC neurons we focused on in the present study seem to fit that description [36, 39].

Morphological analyses performed in mice expressing eGFP in GlyT2^+^ glycinergic neurons revealed that most of these small diameter neurons are in the brainstem, intermingled with giant neurons in the PnC and the PnO [38, 39]. Notably, the projections of these GlyT2^+^ PnC neurons are distributed similarly in the thalamus of mice and man [55]. Previous experiments in rat brain slices showed that PnC glycinergic interneurons are not activated by the stimulation of afferent sensory fibers within the primary startle pathway [40]. Instead, it was suggested that glycinergic interneurons and glycinergic fibers present within the PnC are most likely under the control of excitatory and inhibitory projections from the midbrain and higher brain structures that modulate the startle responses. Here, our morphological reconstruction data (Fig. 7) provide evidence that the amygdala is one of the brain regions that can activate GlyT2^+^ PnC neurons. Since PnC giant neurons in rodents [40, 56] and humans [57] strongly express glycine receptors, we speculate that once activated by the CeA, GlyT2^+^ neurons reduce the excitability of downstream neurons important for PPI including (but not restricted to) giant PnC neurons.

### Behavioral studies

The acoustic startle response can be modulated in various ways. In our current study, the acoustic startle response is *decreased* by a non-startling stimulus preceding the startle stimulus, resulting in a PPI effect. To better understand the different mechanisms involved in modulating the acoustic startle response, it is important to characterize the inputs to PnC neurons at the core of the primary startle pathway and their sensorimotor effects. Recently, functional imaging studies and c-Fos expression data in rats provided strong evidence that CeA neuronal activity is increased during PPI [37]. The objective of our behavioral experiments was to demonstrate the contribution of an CeA-PnC excitatory pathway in reducing the acoustic startle reflex, and its relevance to PPI. We hypothesized that if CeA inputs to the PnC modulate startle during PPI, they would need to be activated *prior* to a startle response. We validated our hypothesis *in vivo* and we showed that unilateral [11, 58–61] photo-inhibition of the CeA-PnC glutamatergic connection during the interval between the prepulse and the pulse decreases PPI (Fig. 5). Interestingly, photo-inhibition of the CeA-PnC glutamatergic connection did not modify an ongoing startle response elicited by a pulse alone startling stimulation (i.e., in the absence of a prepulse). Thus, our data suggest that in the context of PPI, silencing CeA glutamatergic neurons during ISIs between 30-300ms reduces PPI.

During conditioned and unconditioned states of fear, CeA stimulation can *amplify* the acoustic startle response [33–35]. Interestingly, our results showing the role of the CeA during PPI are consistent with the modulatory role of the CeA in fear studies demonstrating that lesions of the CeA block fear *potentiation* of startle without blocking the acoustic startle response itself [46].

In an attempt to mimic an acoustic prepulse and further demonstrate the behavioral relevance of this pathway, we photo-activated CeA-PnC glutamatergic connection prior to an acoustic startling stimulation (Fig. 6). This led to a lower “PPI-like” effect than using an acoustic prepulse. The fact that photo-activation of CeA glutamatergic fibers partially mirrored the effects of acoustic prepulses suggest that CeA excitatory inputs to the PnC do not regulate PPI alone, but work alongside other neuronal elements and pathways, including PPTg-dependent mechanisms. It should also be mentioned that, under physiological conditions, CeA neuronal firing might not necessarily follow the stimulation frequency we used. Despite this, our results clearly show that CeA-PnC excitatory inputs contribute to PPI. Another concern is the possibility that optogenetic inhibition at presynaptic terminals produces unwanted technical effects including a paradoxical increase in neurotransmitter release. To avoid this possibility, we used precisely timed, repetitive light stimulation instead of sustained archaerhodopsin and halorhodopsin activation, which previously led to an increase in spontaneous release [62], allowing us to investigate *in vivo* physiological conditions more closely.

Aside from their role in fear-potentiated startle, no study has directly investigated the function of CeA glutamatergic neurons projecting to the PnC, including GlyT2^+^ neurons. At the level of the PnO, CeA glutamatergic descending projections were suggested to contribute to rapid eye movement (REM)/ active sleep by activating large (i.e., cell body of ∼30um), presumably glutamatergic “REM-on” giant neurons [63]. Since CeA glutamatergic projections target both giant neurons [32] and GlyT2^+^ neurons (present study) in the PnC as well as giant neurons in the PnO [63], it is likely that these projections also target GlyT2^+^ neurons in the PnO. Interestingly, the activation of these two reticular cell types (i.e., giant neurons and GlyT2^+^ cells) leads to opposite behavioral outcomes. That is, in the PnO, giant neurons contribute to REM sleep and GlyT2^+^ neurons are crucial for awake cortical activity [55]. In the PnC, giant neurons are responsible for startle responses [36], and GlyT2^+^ neurons are crucial for behavioral arrest [55]. Our data show that silencing GlyT2^+^ neurons significantly reduced PPI at short interstimulus intervals between the prepulse and the pulse. Based on our data and that of other groups, GlyT2^+^ PnC neurons activated by CeA glutamatergic neurons are likely the ones that inhibit startle during PPI. Future experiments should be done to identify the post-synaptic targets of GlyT2^+^ PnC neurons that are involved in PPI. The amygdala, including the CeA comprises a wide array of molecularly, electrophysiologically, and functionally distinct cell populations (McCullough et al., 2018) that were shown to play differential roles in fear and extinction learning. Therefore, it is possible that a subset of CeA glutamatergic neurons activates a group of lateroventral GlyT2^+^ PnC neurons, leading to PPI, whereas another subset of CeA glutamatergic neurons activates giant PnC neurons, leading to enhanced fear (see schematic of Fig. 7A). These different populations of CeA glutamatergic neurons would likely be activated by inputs from the fear circuit afferent pathways (eg., Hartley et al., 2019) or the PPI circuit afferent pathways.

Amygdalar dysfunctions alter PPI, and both amygdalar function and PPI are abnormal in schizophrenia and other neurological diseases [1, 3, 12, 22, 26, 64–66]. In fact, the CeA also contributes to latent inhibition, a schizophrenia-relevant behavioral assay that measures our ability to ignore/suppress irrelevant stimuli during associative learning. Previously, the effect of the antipsychotic blonanserin was tested in rats where latent inhibition (induced using acoustic stimuli and mild foot shocks) was disrupted by methamphetamine [67]. Disruption of latent inhibition was significantly improved by blonanserin. Immunohistochemical examination showed that blonanserin also increased c-Fos expression in the CeA but not in the basolateral amygdala or the prefrontal cortex. Both PPI and latent inhibition are considered measures of information gating/filtering and their underlying neural circuitry overlap [68]. However, whereas PPI assesses early attentional or “pre-attentive” gating mechanisms, latent inhibition assesses later stages of information gating [69]. Together, these results, along with our *in vivo* findings, suggest that a subset of CeA neurons (including glutamatergic cells) are specifically involved in filtering/inhibiting irrelevant sensory information processing.

### Electrophysiological studies

Previous studies aiming to describe the electrophysiological effect of activating CeA glutamatergic neurons used electrical stimulating electrodes, making it impossible to distinguish whether fibers of passage or CeA cell bodies were activated. Moreover, pharmacological activation of CeA neurons did not allow to selectively activate excitatory neurons [32]. To reconcile the glutamatergic nature of the CeA neurons projecting to the PnC with their contribution to an inhibitory phenomenon, here, we performed *in vitro* patch clamp recordings in GlyT2-CRE mice (Fig. 8). We recorded EPSPs in GlyT2^+^ PnC neurons labeled with tdTomato, in response to targeted activation of CeA glutamatergic inputs expressing ChR2. Altogether, our results demonstrate that: 1-CeA glutamatergic projections activate GlyT2^+^ PnC neurons via AMPA and NMDA receptors, and 2-CeA-PnC glutamatergic synapses display short-term synaptic facilitation.

Previous rat pharmacological *in vivo* studies showed that blocking AMPA and NMDA receptors in the PnC inhibits acoustic startle responses [70–72]. Interestingly, our behavioral data suggest that blocking a glutamatergic connection between the CeA and the PnC reduces PPI. Our PPI *in vivo* results do not conflict with these previous studies showing the important role of the glutamate neurotransmission within the PnC startle pathway. Our *in vivo* results clearly indicate that during PPI, inhibition of startle involves CeA glutamatergic cells directly projecting to the PnC and activated by prepulses. In that context, acoustic prepulses activate glutamatergic receptors located on GlyT2^+^ and likely also, PnC giant neurons. One hypothesis is that during PPI, CeA glutamate neurotransmission activates GlyT2^+^ PnC neurons leading to a feedforward inhibition. This hypothesis is clearly supported by our *in vitro* electrophysiological recordings in PnC slices showing that AMPA/NMDA receptor blockers reduce the EPSPs in GlyT2^+^ PnC neurons elicited by CeA glutamatergic fiber stimulation (Fig. 8C).

Paired photo-stimulation at CeA-PnC glutamatergic synapses elicited short-term synaptic facilitation of EPSPs at ISI of 50 and 100ms (Fig. 8B). Synaptic facilitation reflects presynaptic enhancement of neurotransmitter release, associated with residual calcium accumulation within presynaptic terminals following the first stimulation [73]. Interestingly, the synaptic facilitation we recorded at CeA-PnC glutamatergic synapses and the CeA-dependent PPI values we measured *in vivo* both occur within similar time scales.

Typically considered as a GABAergic (and non-cholinergic) nucleus, the CeA includes an ensemble of several other neurochemical and neuropeptide profiles, such as galanin, somatostatin, substance P, corticotropin-releasing factor [76–80]. In fact, the CeA sends GABAergic inputs to several brainstem regions adjacent to the PnC, such as the ventrolateral periaqueductal grey [81], locus coeruleus [82] and nucleus of the solitary tract [83]. Furthermore, altered GABAergic neurotransmission is seen in schizophrenia [84, 85] and is associated with abnormal acoustic startle reflex and PPI [86–90]. GABAergic neurotransmission was previously shown to modulate the startle pathway [91]. Although our data rule out the possibility that the CamKIIα^+^ CeA neurons we targeted are inhibitory (Figs. 4 and 8), future work should determine whether the CeA also contribute to PPI through GABAergic projections. Previous studies have highlighted the importance of pontine glutamatergic and GABAergic signaling during PPI. In addition, PPI is sensitive to changes in glutamatergic and GABAergic transmission in several brain regions, such as the amygdala [26, 27, 86], hippocampus [87], superior colliculus [88], PPTg [20, 21, 89] and nucleus accumbens [90]. These brain regions also exhibit anatomical and functional abnormalities in neuropsychiatric disorders associated with sensorimotor gating deficits. Here, we provide evidence for a potential amygdala-dependent glutamatergic mechanism at the PnC level, that could be impaired in diseases associated with PPI deficits.

## Conclusions

Overall, the results presented here along with the body of literature on the role of the amygdala on acoustic startle modulation, suggest that CeA excitatory neurons send inputs to the PnC, where they activate a subpopulation of GlyT2^+^ neurons that contribute to PPI. We propose that the primary startle pathway (Fig. 1; red), is modulated by two parallel circuits: **1-** the well-established the CSPP network (Fig. 1; light blue and dark blue brain regions/pathways), and **2-** the CeA-PnC glutamatergic connection (Fig. 1; dark blue dotted connection delineated by the red dotted square). These two circuits ultimately converge at the level of the PnC. Our results are aligned with recent data obtained by other groups revisiting the theoretical constructs of PPI, describing glutamatergic, gabaergic and glycinergic underlying mechanisms. More importantly, our results shed light into the basic processes underlying sensory filtering, and our disease-relevant proposed circuit should expand insights derived from disease experimental systems.

## Materials and Methods

### Mice

Experiments were performed on C57BL/6 male mice (N=61; The Jackson Laboratory, Bar Harbor, ME), GlyT2-eGFP mice (N=6; graciously provided by Dr. Manuel Miranda-Arango, University of Texas at El Paso, El Paso, TX) and GlyT2-Cre^+/-^ mice (N=10; graciously provided by Dr. Jack Feldman, University of California, Los Angeles). Litters were weaned at PND 21 and housed together until stereotaxic microinjections were performed at PND 70-84 (adult). Mice received food and water *ad libitum* in a 12hour light/dark cycle from 7:00 am to 7:00 pm. This age corresponds to the age of the animals used in the Paxinos and Franklin Mouse Brain Atlas, from which all the stereotaxic coordinates were derived, and cytoarchitectural boundaries delineated [42]. Following surgical procedures, mice were single-housed and monitored for the duration of the recovery period. Experiments were performed in accordance with and approved by the Institutional Animal Care and Use Committee of the University of Texas at El Paso (UTEP) and the University of Massachusetts Amherst (UMass).

### Stereotaxic microinjections

Mice were sedated by inhaling 5% isoflurane vapors (Piramal Critical Care, Bethelehem, PA), then placed on a stereotaxic apparatus (model 900, David Kopf, Tujunga, CA) and immobilized using ear bars and a nose cone. Mice were maintained under 1.5%-2% isoflurane throughout the duration of the surgical procedure. With bregma as a reference, the head of the mice were leveled on all 3 axes. A craniotomy was performed directly dorsal to the injection site. Then, using a microinjector (Stoelting Co., Wood Lane, IL) with a 5μl Hamilton syringe (Hamilton Company Inc., Reno, NV) and a 32 gauge steel needle, unilateral injections of 50-80nl of the retrograde neuronal tracer Fluoro-Gold (Molecular Probes, Eugene, OR, catalog# H22845, lot# 1611168) were infused into the PnC (coordinates from bregma: AP −5.35mm; ML +0.5mm, DV −5.6mm; N=4 mice). The CAG-FLEX-tdTomato or rAAVDJ/Ef1α-DIO-eArch3.0-eYFP viral vectors were injected (200nl) in the PnC of mice expressing the CRE recombinase enzyme in GlyT2^+^ neurons (GlyT2-Cre mice; N=10). In separate animal cohorts, 100-125nl of AAV particles were unilaterally injected in the CeA (AP −1.35mm, ML +2.66mm, DV −4.6mm). For these viral injections, pAAV DJ-CamKIIα-eArch3.0-eYFP, pAAV DJ-CamKIIα-NpHR3.0-eYFP, pAAV DJ-CamKIIα-hChR2(H134R)-eYFP, pAAV DJ-CamKIIα-eYFP or pAAV DJ-CamKIIα-mCherry viral particles were used (4 × 10^12^ particles/mL; all vectors were obtained from the Deisseroth Lab/Optogenetics Innovation Lab, Stanford University, Palo Alto, CA). Fluoro-Gold and viral particles were delivered at a rate of 50nL/min. The microinjection syringe was left in place for 10 minutes after infusion to limit spillover during needle retraction. Mice injected with Fluoro-Gold recovered for 5-7 days, to allow optimal Fluoro-Gold retrograde transport to occur. AAV-injected mice recovered for 3-5 weeks to allow sufficient time for maximal viral transduction.

### Immunohistochemistry

Mice were perfused transcardially with 0.9% saline solution for 10mins followed by 4% paraformaldehyde (PFA) in 0.1M phosphate buffer saline (PBS; pH 7.4) for 15 mins, brains were then extracted and post-fixed overnight in 12% sucrose in PFA solution. After three 0.1M PBS rinses (5 minutes each), brains were frozen in chilled hexanes for 1 minute and stored at −80°C. Using a microtome, four 1-in-5 series of 30μm coronal sections were cut, and stored in cryoprotectant (50% 0.05 M phosphate buffer, 30% ethylene glycol, 20% glycerol) at −20°C. One of the series was rinsed three times (5mins each) with 0.1M Tris-buffered saline (TBS; pH 7.4), mounted and coverslipped to visualize injection and projection sites. An adjacent series of brain sections was Nissl-stained to determine plane of section, and delineate cytoarchitectural boundaries. The two remaining series were used for immunohistochemistry. For mice injected with Fluoro-Gold, coronal tissue sections at the level of the PnC, CeA and PPTg were washed with 0.1M TBS (5 washes, 5 mins each), and incubated in blocking solution (2% normal donkey serum, 0.1% Triton X-100; in 0.1M TBS) for 1-2 hours at room temperature. PPTg sections were incubated with a goat anti-ChAT primary antibody (1:100, Millipore, catalog# AB144P-200UL, lot# 2854034) for 60 hours at 4°C, washed with TBS, and then incubated in a Cy3-conjugated donkey anti-goat secondary antibody (1:500, Jackson ImmunoResearch Laboratories, catalog# 705-165-147, lot# 115611) for 4-5 hours at room temperature. Tissue slices were then washed with TBS, mounted and coverslipped. Similarly, for mice injected with viral particles, tissue sections containing the PnC, CeA and the PPTg were incubated in a chicken anti-GFP primary antibody (1:1000, Abcam, catalog# ab13970, lot# GR236651-13), then incubated in an Alexa Fluor 488-conjugated donkey anti-chicken secondary antibody (1:500, Jackson ImmunoResearch Laboratories, catalog# 703-545-155, lot# 130357), followed by incubation in NeuroTrace™ (640/660 deep red fluorescent nissl stain, 1:100 in TBS, ThermoFisher, catalog# N21483). NeuroTrace™ was alternatively used to determine plane of section and cytoarchitecture. Tissue sections at the level of the PnC of GlyT2-eGFP mice injected with pAAV DJ-CamKIIα-mCherry in the CeA (N=6 mice) were incubated with a chicken anti-mCherry (1:1000, Abcam, catalog# ab205402, lot# GR225123-3) and a rabbit anti-PSD95 (1:500, Abcam, catalog# ab12093, lot# GR317630-1) primary antibodies. Then, sections were incubated with a Cy3-conjugated donkey anti-chicken (1:500, Jackson ImmunoResearch Laboratories, catalog# 703-165-155, lot# 130328) and a Cy5-conjugated donkey anti-rabbit (1:500, Jackson-ImmunoResearch Laboratories, catalog# 705-545-147, lot#125100) as described above.

### *In situ* hybridization

Mice injected with pAAVDJ-CamKIIα-eYFP in the CeA (N=3 mice) were anesthetized with inhaled isoflurane and rapidly decapitated. Brains were harvested, frozen in chilled isopentane and stored at −80°C. Serial coronal sections (15µm) at the level of the CeA were cut in a cryostat, directly mounted onto glass slides, and stored at −80°C. Tissue sections on slides were submerged in freshly prepared cold 4% PFA for 15 mins, rinsed twice briefly with 0.1M Phosphate Buffer (PB) and dehydrated in increasing ethanol solutions (50%, 70%, 100%, 100%; 5 mins each at room temperature). Then, the RNAscope assay (Advanced Cell Diagnostics) started by incubating in hydrogen peroxide (H_2_O_2_) for 10 mins in a humidified box, followed by protease III incubation for 15 mins. RNA hybridization probes against genes encoding mouse VGLUT2 (319171-C1), and eYFP (312131-C2) were then incubated for 2hr at 40°C. Antisense probes were also included as controls in a separate glass slide. Probe signals were then developed separately with Opal Dyes (opal 690 1:1.5K, opal 520 1:750), and coverslipped with ProLong Gold™ with DAPI.

### Morphological Reconstruction

Z-stacks from tissue sections of GlyT2-eGFP and GlyT2-Cre mice were obtained on a Nikon A1 Resonant Confocal microscope (Nikon Instruments Inc., Melville, NY) equipped with NIS-Elements High Content Analysis software (version 5.02, Nikon Instruments Inc., Melville, NY). Tissue sections containing labeled CeA and PnC neurons were first examined on a single Z-plane with the 10x objective to survey the tissue section. Using a 60x objective, an area (212.56µm width x 212.56µm height) within CeA and PnC sections were then sequentially scanned by the 488, 561 and 640nm laser lines in 0.1µm Z-steps throughout the 30µm tissue section. Z-stacks were analyzed with NIS-Elements 5.0 Advanced Research software (version 5.02, Nikon Instruments Inc., Melville, NY). To visualize close appositions of CeA axons (labeled with mCherry) with GlyT2^+^ neurons (labeled with eGFP) in GlyT2-eGFP mice, a binary layer was configured to segregate putative synaptic contacts of >50nm in distance (due to technical limitations). These contacts were imaged in split-channels and orthogonal views. Then, z-stacks were reconstructed in three-dimension and volume was rendered.

### Nissl stain

Series of tissue slices were mounted on gelatin-coated slides and air-dried overnight. Slides were immersed in deionized water, followed by ascending concentrations of ethanol (3mins each: 50%, 75%, 95% and 100%), and then in xylenes (30mins). Brains slices were rehydrated in descending concentrations of ethanol and DI water, dipped 12-20 times in a thionin acetate solution, and then washed in DI water. Brains slices were dehydrated, slides were then coverslipped with DPX, and air-dried overnight.

### Microscopy analysis

Tissue sections were analyzed with an Axio Observer.Z1 epifluorescence microscope (Carl Zeiss Inc., Thornwood, NY) equipped with Fluoro-Gold, GFP, Cy3 filters, 10x and 40x objectives, a motorized stage and Axiovision Rel. 4.8 software (Carl Zeiss Inc., Thornwood, NY). To create photomontages, single Z-plane images were obtained with the MosaiX module of the Axiovision Rel. 4.8 software at 10x for each fluorophore sequentially (1024 × 1024 pixel resolution). Images acquired for the intensity and quantification of eYFP fluorescence analysis were captured and processed using identical settings. A total of 836 images (fluorescence and bright-field) were analyzed for each brain region. Nissl stained slices were imaged using bright field microscopy, and boundaries were delineated using Adobe Illustrator (Adobe, San Jose, CA).

### Electrophysiological recordings

Whole-cell recordings were performed using glass pipettes (3–5 MΩ) filled with intracellular solution (in mM): KMeSO4 (125), KCl (10), HEPES (10), NaCl (4), EGTA (0.1), MgATP (4), Na2GTP (0.3), Phosphocreatine (10), Biocytin (0.1%) (pH=7.3; osmolarity=285–300 mosm). The glass microelectrode was mounted on a patch clamp headstage (Molecular Devices LLC, Sunnyvale, CA; catalog# CV-7B), which was attached to a multi-micromanipulator (Sutter Instrument, Novato, CA; catalog# MPC-200). Data were acquired with pClamp10 software using a MultiClamp™ 700B amplifier (Molecular Devices LLC, Sunnyvale, CA) and a Digidata 1550B digitizer (Molecular Devices LLC, Sunnyvale, CA). EYFP-expressing CeA cells and tdTomato-expressing GlyT2^+^PnC cells were imaged and targeted using NIS-Elements Basic Research software (version 5.11, Nikon Instruments Inc., Melville, NY). Only cells with an initial seal resistance greater than 1GΩ, a resting membrane potential between −60mV and −70mV, a holding current within −100pA to 100pA at resting membrane potential and overshooting action potentials were used.

In CeA slices, 15 pA depolarizing current steps were injected for 500 ms to induce action potentials in CeA neurons expressing CamKIIα-ChR2-eYFP, in current clamp. Spontaneous EPSCs were recorded at a holding potential of −70 mV, in voltage clamp. Evoked EPSPs were also recorded in these CeA neurons held at −70mV, in response to a 1-ms blue light pulse. Blue light was delivered every 30 secs using a 200μm optic fiber mounted on a micromanipulator connected to a blue LED (473nm; Plexon, Dallas, TX) and positioned in close proximity to the recorded neuron.

In PnC slices, electrical properties of the GlyT2^+^ neurons were first recorded in voltage clamp. Spontaneous excitatory postsynaptic currents (sEPSC) were recorded for 5 mins at −70 mV, and inhibitory postsynaptic currents (IPSCs) were recorded for 5mins at 0mV. Then, in current clamp mode, 15 pA depolarizing current steps (from −150pA to 150pA) were injected for 500 ms to analyze the spiking properties of GlyT2^+^ cells. Pulses of blue light (1 ms), applied every 30secs, were used to photo-stimulate CeA excitatory fibers in PnC slices. The photo-stimulation elicited EPSPs and EPSCs in GlyT2^+^ neurons, held at −70mV. Paired light pulses with 50 and 100ms ISI were also delivered to characterize short-term plasticity. GlyT2^+^ neurons were then held at 0mV, to record light-evoked inhibitory post-synaptic current (IPSCs) or potentials (IPSPs). The NMDA receptor antagonist AP5 (50 μM) and the AMPA receptor antagonist DNQX (25 μM), were freshly diluted prior to use. At synapses between CeA excitatory cells and GlyT2^+^ neurons, synaptic events were recorded for 10 minutes in aCSF. Then, 20 mins after the bath application of glutamate receptors antagonists, synaptic events were recorded during 10-minutes in the presence of the antagonists. This was followed by a 20 minutes washout period, and synaptic events were recorded during the following 10 mins, in aCSF.

At the end of all whole cell recordings, the cell membrane was sealed by forming an outside-out patch. The glass microelectrode was slowly retracted, and as the series resistance increased, the membrane potential was clamped at −40mV. The 300μm-thick acute brain slices containing the recorded cells (CeA or PnC) were immersed in 4% PFA solution overnight. Following overnight PFA fixation, these brain slices were rinsed with PBS (3 times, 5mins each). Slices were then incubated in anti-RFP and/or anti-GFP antibodies and in complementary secondary antibodies to enhance the fluorescence of the viral vectors used. Following PBS rinses, slices were incubated with Cy5-conjugated streptavidin (a biotin-binding protein) diluted in PBS (with 0.1% Triton X-100) at room temperature for 4-5 hours or overnight at 4°C. Slices were then rinsed with PBS, mounted on glass slides, coverslipped and sealed with ProLong™ Gold antifade reagent (Invitrogen by ThermoFisher Scientific, Waltham, MA, catalog# P36934, lot# 1943081), and air dried overnight in the dark.

### Behavioral testing

Three to four weeks after the viral injection in the CeA, non-injected WT control mice and WT mice injected with a viral vector were sedated by inhaling 5% isoflurane vapors, placed in a stereotaxic apparatus, and immobilized using ear bars and a nose cone. Mice were maintained under anesthesia (1.5%-2% isoflurane), the head was leveled in all three axes. With bregma as a reference, a craniotomy was drilled directly dorsal to the implantation site, at the PnC level. A cannula guide with a 200μm core optical fiber (Thorlabs, Newton, NJ) was then implanted over the PnC (AP −5.35mm, ML +0.5mm, DV −5.3mm), and cemented to the skull with dental cement (Parkell, Edgewood, NY). Mice recovered for 7 days post-surgery before behavioral testing. Mice underwent the PPI task in a startle response system (PanLab System, Harvard Apparatus, Holliston, MA). Behavioral testing trials were designed, and data were recorded using PACKWIN V2.0 software (Harvard Apparatus, Holliston, MA). Sound pressure levels were calibrated using a standard SPL meter (model 407730, Extech, Nashua, NH). Mice were placed on a movement-sensitive platform. Vertical displacements of the platform induced by startle responses were converted into a voltage trace by a piezoelectric transducer located underneath the platform. Startle amplitude was measured as the peak to peak maximum startle magnitude of the signal measured during a 1s window following the presentation of the acoustic stimulation. Prior to any testing session, animals were first handled and acclimatized to the testing chamber, where the mice were presented to a 65dB background noise, for 10 mins. This acclimatization period was used to reduce the occurrence of movement and artifacts throughout testing trials. Following the acclimatization period, an input/output (I/O) assay was performed to test startle reactivity. This I/O test began with the presentation of a 40-ms sound at different intensities (in dB: 70, 80, 90, 100, 110 and 120) every 15s, in a pseudorandomized order. Background noise (65dB) was presented during the 15s between sounds. A total of 35 trials (i.e. 7 sound intensities, each sound presented 5 times) were acquired and quantified. Startle reactivity, derived from this I/O assay, allowed the gain of the movement-sensitive platform to be set. This gain allowed the startle responses to be detected within a measurable range. Once determined, the gain for each experimental subject was kept constant throughout the remaining of the experiment. Following a one-hour resting period, mice were presented with seven startle-inducing 120 dB (40 ms) sounds called “Pulse-alone” stimulations. These 120 dB sounds were presented every 29 secs (interspersed with 65dB background noise), and were used to achieve a stable baseline startle response. The following PPI test consisted of two different conditions as follows: 1- startling pulse-alone stimulations (for baseline startle amplitude), and 2- combinations of a prepulse (75 dB noise; 20 ms) followed a 120 dB startling pulse (40 ms) at 8 different interstimulus intervals (in ms): 10, 30, 50, 100, 200, 300, 500, and 1000 (end of prepulse to onset of startle pulse). The inter-trial interval of these two conditions was 29 secs.

For combined optogenetic manipulations, animals injected with either control viral vectors or vectors containing ChR2 (N=8 mice), Arch3.0 (N=8 mice), NpHR3.0 (N=8 mice) or the control vector (pAAV DJ-CamKIIα-eYFP; N=8) were tested in the startle chamber. These animals were closely monitored to ensure that they were comfortably tethered to an optic fiber, which exited through a small opening from the roof of the startle chamber. The optic fiber (200 µm diameter, Thorlabs, Newton, NJ) was connected to the animal’s head via a cannula implanted on the head of the mouse with a zirconia sleeve (Thorlabs, Newton, NJ). Animals were tethered ∼15 mins before testing and allowed to move freely, exploring their home cages before being transferred to the startle chamber. Optogenetic stimulation was triggered by a signal from the Packwin software (PanLab System; Harvard Apparatus, Holliston, MA), which was transformed into a TTL-pulse. This TTL-pulse triggered a waveform generator (DG1022, Rigol Technologies), which was used to modulate light stimulation. Photo-stimulation was delivered using a blue 473nm laser (Opto Engine LLC, Midvale, UT) for ChR2 activation. Photo-inhibition was delivered using a yellow 593.5nm laser (Opto Engine LLC, Midvale, UT) for NpHR3.0 activation, or a green 532nm LED (Plexon, Dallas, TX) for Arch3.0 activation. During PPI trials paired with optogenetic manipulations, a train of light stimulation (1ms light ON, 200ms light OFF) was delivered at 5Hz and was either: 1- paired with the pulse-alone stimulation, or 2- delivered 1sec before the prepulse, lasting the entire ISI. During PPI trials with optical stimulation used as a prepulse, a 5Hz or 20Hz stimulation train (3 pulses of 15ms) was delivered at various ISI (10, 30, 50, 100, 200, 300, 500, and 1000 ms; end of prepulse to onset of startle pulse) prior to the startling pulse. At the end of each experiment, histological analyses were performed to confirm that: 1- the injected viral particles were confined to the CeA, and 2- the cannula guide placement was successfully aimed at the PnC. If these criteria were not met, the subject was excluded from the study.

### Statistical analysis

Cell counting of eYFP-labeled or VGLUT2-expressing somata within the CeA was performed in a tissue slice series of 6 slices spanning levels 40 to 44 of the Paxinos and Franklin Mouse Brain Atlas [42]. Imaging was performed as outlined in the Microscopy Analysis section. Percentages of labeled somata were calculated as eYFP+/Neurotrace-labeled cells (Fig 3) or VGLUT2+/eYFP+ (Fig 4). Statistical analyses were performed using SigmaPlot (Systat Software, Inc., San Jose, CA). Normality and equal variance of the data were first tested, and data transformations were made before performing further statistical analyses. We determined the significance of the interaction between the factors assessed using ANOVA. For the results of whole-cell patch-clamp recordings with receptor antagonists, one-way repeated-measures (RM) ANOVA and Tukey post-hoc testing were used to assess the effect of the receptor antagonists on the light-evoked events. For PPI in vitro results, one-way ANOVAs and Tukey post-hoc testing were used to reveal if at any ISI the electrically-evoked fEPSPs were significantly attenuated by the optical stimulation of CeA-PnC excitatory synapses. PPI was defined and measured as [1–(startle amplitude during “Prepulse+Pulse” trials/startle amplitude during “Pulse” trials)] × 100. Two-way RM ANOVA was used to assess the effect of the vector used, light, sound intensity/ISI and light interaction and the interaction among groups. Then Tukey testing was applied for post-hoc comparisons. For optical stimulation experiments where the photo-stimulation of CeA fibers was used as a prepulse *in vivo*, two-way RM ANOVA was used to assess the effect of the stimulation modality/frequency used, ISI, ISI and stimulation modality/frequency interaction and the interaction among groups. Then, Tukey testing was applied for post-hoc comparisons. A confidence level of p<0.05 was considered statistically significant. Sample sizes were chosen based on expected outcomes, variances and power analysis. Data are presented as means ± SEM. N indicates total number of animals; n indicates total number of brain slices or testing trials. Adobe Illustrator was used to create figures.

## Supporting information

Cano et al., 2021 Supplemental File

## Declarations

### Availability of data and materials

The datasets used and/or analyzed during the current study are available from the corresponding author on reasonable request. All data generated or analyzed during this study are included in this published article and its supplementary information files. Details about the GlyT2-Cre mice can be found in Giber et al., 2015 [55]. Details about the GlyT2-eGFP mice can be found in Zeilhofer et al., 2005 [39].

### Authors’ contributions

J.C.C. and K.F. conceptualized the study; J.C.C. and W.H., performed the experiments, analyzed the data and perform the statistical analyses; J.C.C. and K.F. wrote the paper; J.C.C., W.H. and K.F. revised the final version of the paper. All authors read and approved the final manuscript.

## Acknowledgments

We thank Ms. Carla D. Loyola (UTEP) for the technical support and for the animal care; Dr. Stephanie Padilla (UMass), Dr. Jack Feldman (UCLA), Dr. Manuel Miranda-Arango, Dr. Ellen M. Walker and Ken Negishi (UTEP) for the technical assistance; Dr. Joseph A. Gogos (Columbia University), Dr. Amy B. MacDermott (Columbia University), Dr. Arshad M. Khan (UTEP) and Dr. Paul S. Katz (UMass) for the insightful discussions; and Dr. James Chambers for the assistance with the three dimensional reconstructions performed at the Light Microscopy Facility at UMass Amherst.

## Ethics approval and consent to participate

Not applicable.

## Consent for publication

Not applicable.

