## Supplementary material for "Shedding light on the contribution of amygdalar excitatory projections to prepulse inhibition of the auditory startle reflex": Cano et al., 2021 Supplemental File

### Additional File 1

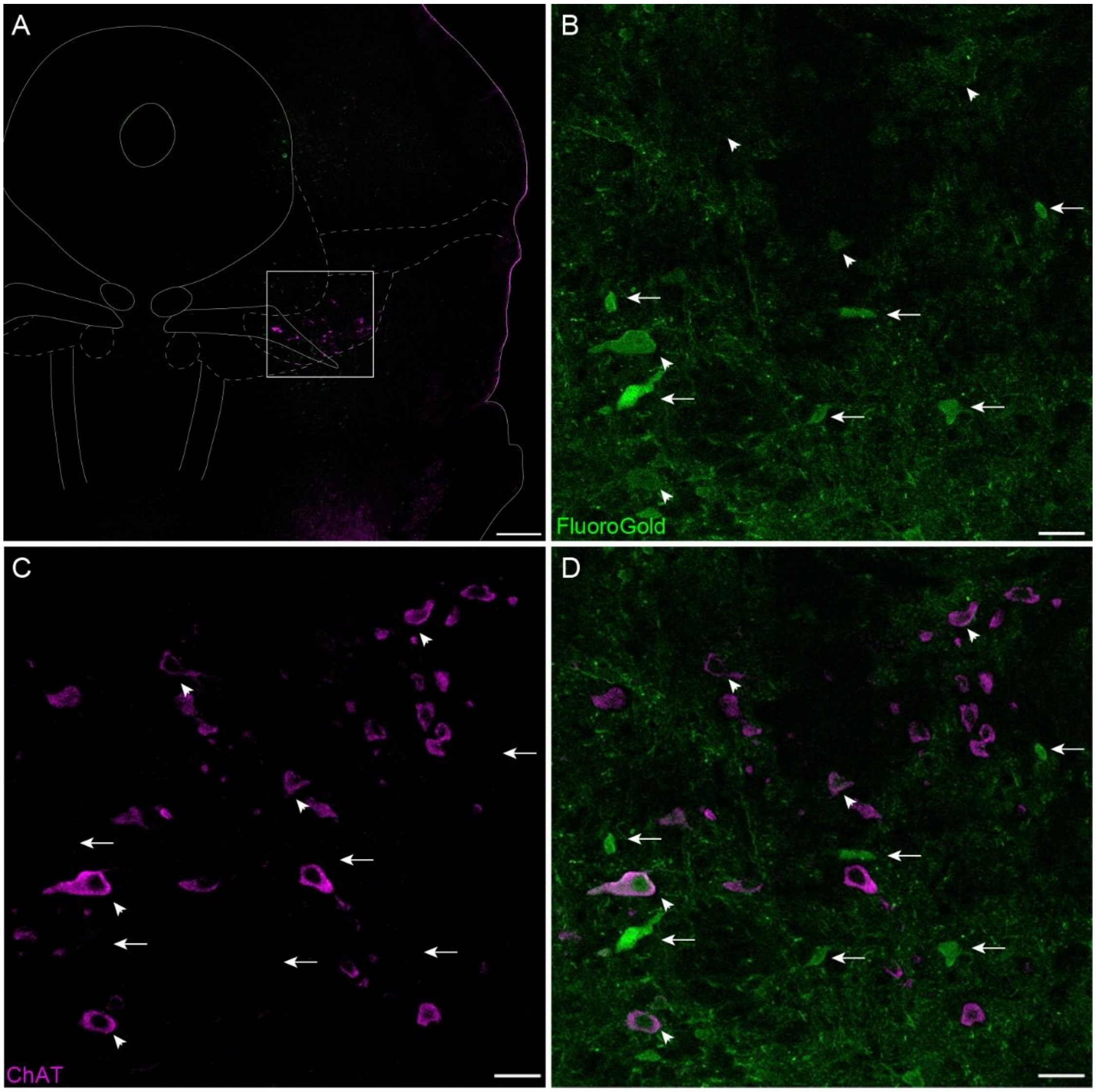


**Supplemental Figure S1. The PnC receives monosynaptic inputs from the PPTg.** (A) Representative PPTg coronal section showing the immunofluorescence of the cholinergic marker ChAT (magenta), which delineates the PPTg. (B) PPTg $ChAT^+$ cell bodies (magenta) shown at higher magnification. (C) Representative PPTg section showing Fluoro-Gold staining (green). (D) Overlay of B and C, representative co-immunostaining of Fluoro-Gold and ChAT fluorescence. Arrows indicate neurons stained with Fluoro-Gold that are non-cholinergic ($ChAT^-$). Arrowheads indicate neurons neurons stained with Fluoro-Gold that are $ChAT^+$. Representative of N = 4 mice. Scale bars: (A) 500µm, (B-D) 100µm.

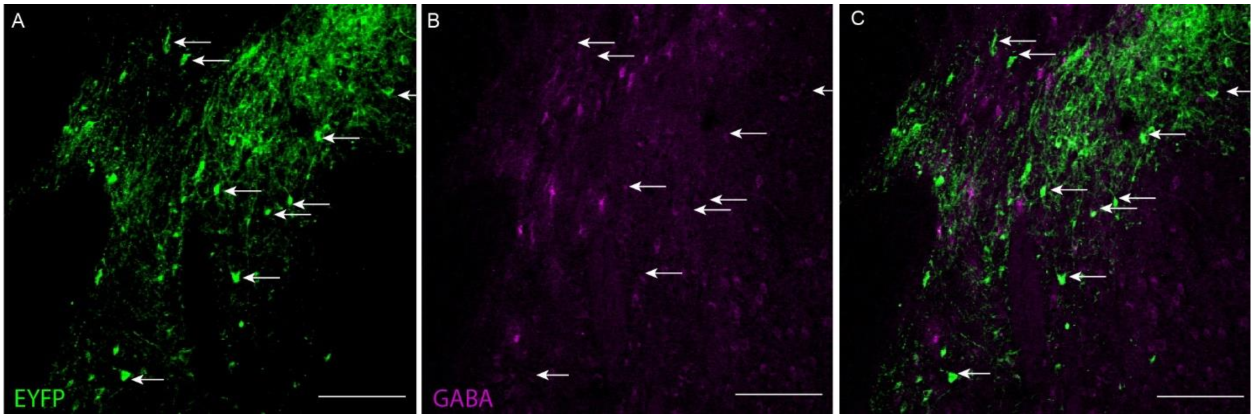


**Supplemental Figure S2. CamKIIα⁺ CeA projection neurons are non-GABAergic.** (A) Representative images of GABA (magenta) immunostaining in the medial division of the CeA. Arrowheads indicate $GABA^{+}$ CeA cell bodies. (B) Representative image of eYFP fluorescence (green). Arrows indicate CeA cell bodies and neurites. (C) Overlay of A and B, representative co-immunostaining for eYFP and GABA showing no overlap. Representative of N = 4 mice. Scale bars: (A-C) 250µm.

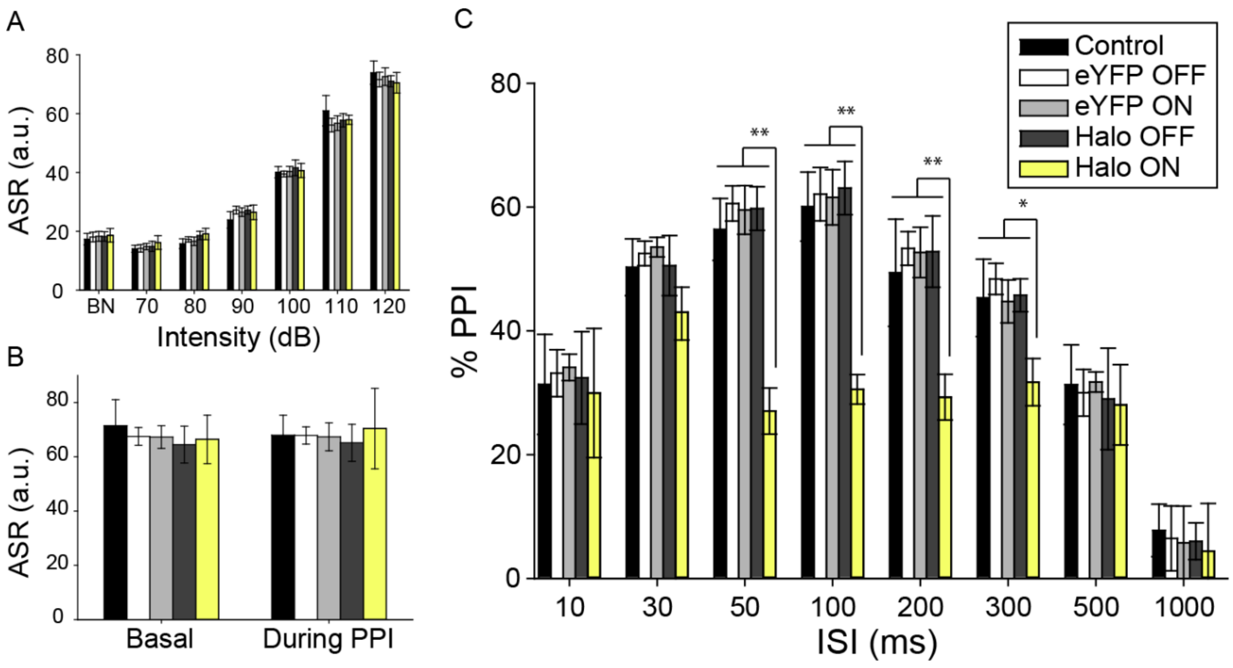


**Supplemental Figure S3. NpHR3.0 inhibition of the CeA-PnC excitatory connection reduces PPI.** (A) Graph showing no significant main effect of yellow light on mean baseline startle amplitude [sound: ($F_{(1,11)}$=1.935, p=0.115); yellow light: ($F_{(1)}$=0.00297, p=0.958); sound intensity*light interaction ($F_{(1,6)}$=0.102, p=0.996)] by comparing non injected WT control mice, mice injected with eYFP only (light ON or OFF) and mice injected with Halorhodopsin (NpHR3.0; light ON or OFF). (B) Graph showing no significant main effect of light during 120dB pulses presented before (basal) vs. randomly during the PPI task, on mean baseline startle amplitude among animal groups ($F_{(1)}$=0.394, p=0.543). (C) Graph showing that optogenetic silencing of CeA-PnC excitatory synapses during prepulses significantly decreased PPI in mice injected with NpHR3.0 at ISIs between 50 and 300ms. We found a significant effect of ISI ($F_{(1,7)}$=33.019, p<0.001), light ($F_{(1)}$=5.371, p=0.041) and the light*ISI interaction ($F_{(1,7)}$=3.692, p=0.002) on PPI (ANOVA). N=8 mice per group. Data are represented as mean ± SEM. *p<0.05, **p<0.01.

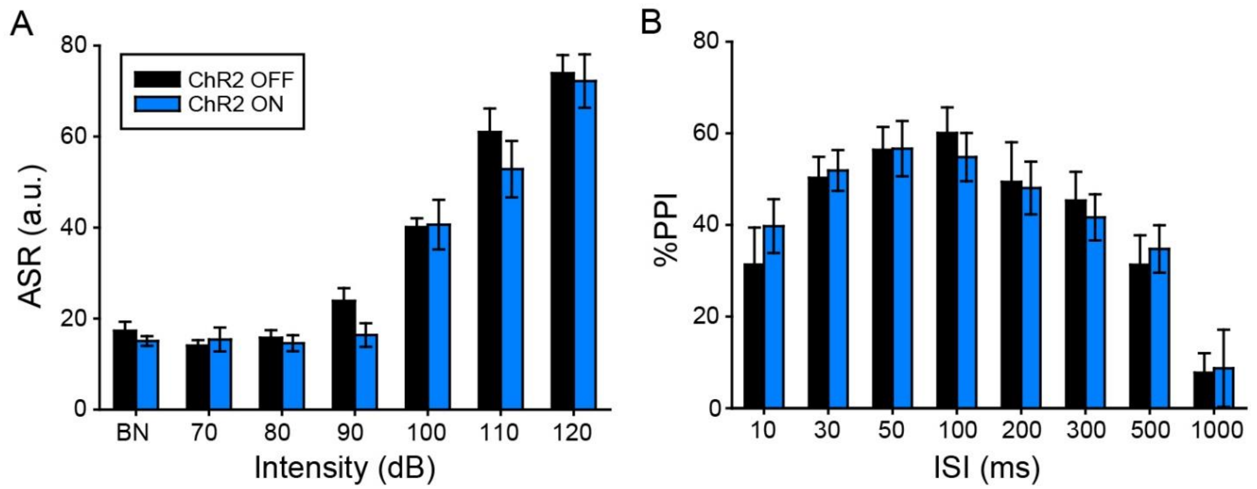


**Supplemental Figure S4. Photo-stimulating CeA-PnC glutamatergic synapses does not alter acoustic PPI.** (A) Graph showing no significant main effect of blue light on mean baseline acoustic startle amplitude in mice injected with ChR2 (light ON or OFF). We found no effect of viral vector type ($F_{(1,2)}$=1.417, p=0.247) or viral vector*sound intensity interaction ($F_{(1,12)}$=0.413, p=0.956). (B) Graph showing no significant main effect of blue light paired with acoustic prepulses on PPI, at all ISIs tested ($F_{(1,14)}$=0.151, p=1.000). N=6 mice. Data are represented as mean ± SEM.

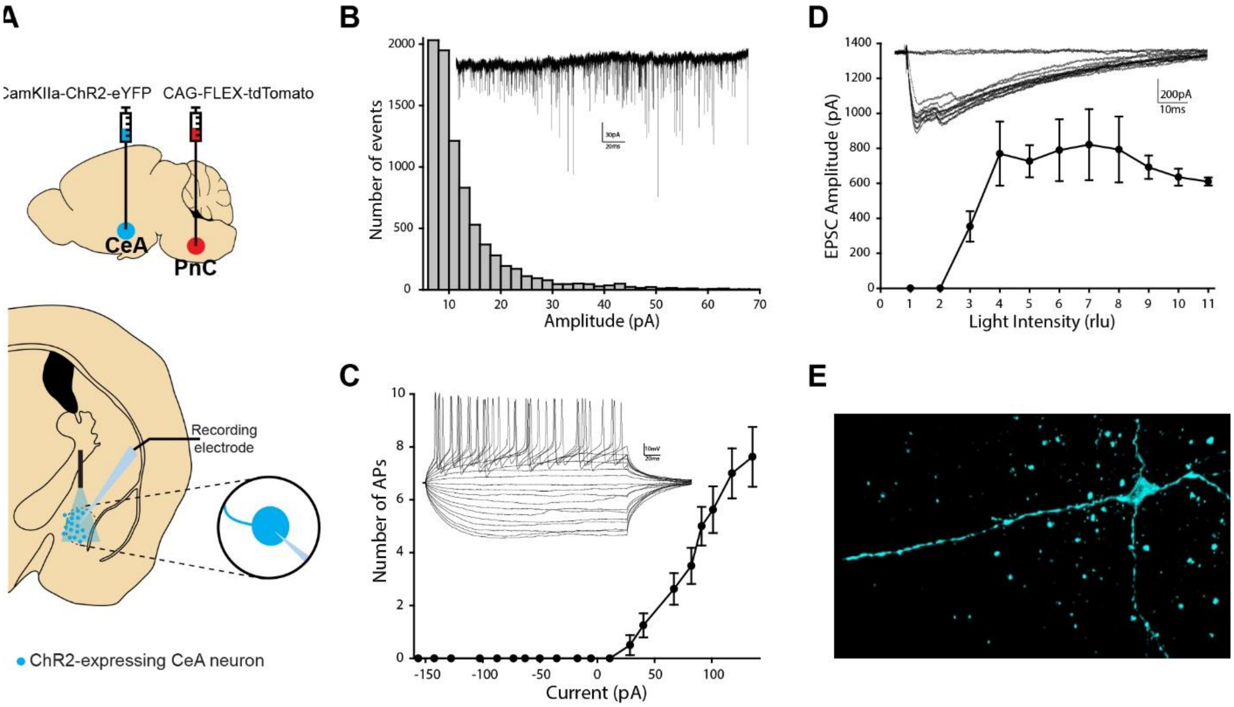


**Supplemental Figure S5. Intrinsic and synaptic properties of CamKIIα-eYFP$^+$ CeA glutamatergic neurons expressing ChR2.** (A) Scheme of the *in vitro* patch clamp recording experiment showing the injection site of AAVDJ-CamKIIα-ChR2-eYFP and the recording of CeA neurons of GlyT2-Cre mice. (B) Plot showing the cumulative distribution of sEPSC amplitude. *Inset*, representative trace. (C) Plot of the firing rate as a function of depolarizing currents. *Inset*, representative traces. (D) Input/output curve of light-evoked EPSCs. (E) Three-dimensional reconstruction representative of a recorded CeA neuron filled with biocytin. N=10 mice, n=26 neurons. Data represented as mean ± SEM. *P>0.05, **P>0.01.

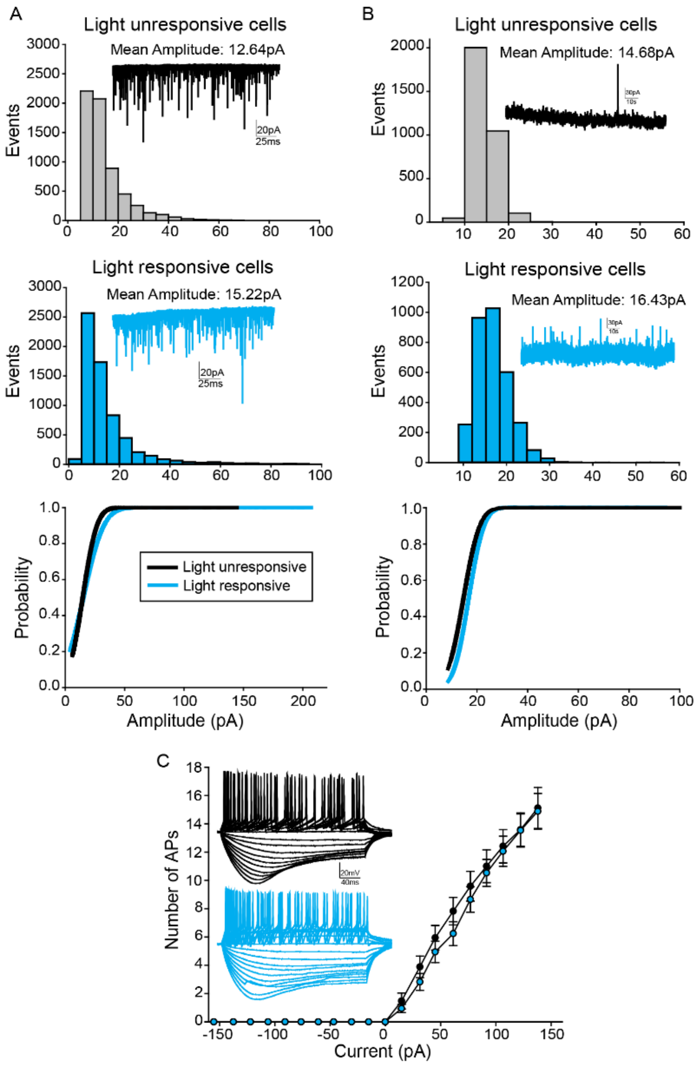


**Supplemental Figure S6. Intrinsic and synaptic properties of GlyT2$^+$ PnC neurons.** (A) Plot showing the cumulative distribution of sEPSC amplitude recorded in GlyT2$^+$ cells unresponsive (*Top*; n=20 cells) and GlyT2$^+$ cells responsive (*Middle*; n=18) to light. *Bottom*: cumulative sEPSC distribution plots. (B) Plot showing the cumulative distribution of sIPSC of GlyT2$^+$ cells unresponsive (*Top*; n=20 cells) and GlyT2$^+$ cells responsive (*Middle*; n=18) to light. *Bottom*: cumulative sIPSC distribution plots. (C) Graph showing no significant difference between the firing rate and threshold current to elicit APs in GlyT2$^+$ cells unresponsive and responsive to light. *Insets*: Representative traces of N=10 mice, n=38 cells. Data represented as mean ± SEM.

| **Group** | **$R_a$** | **$R_m$** | **$C_m$** | **$I_h$** |
|---|---|---|---|---|
| Light unresponsive (n=20) | 65.33 ± 5.69 | 212.42 ± 21.6 | 28.5 ± 4.15 | -8.53 ± 2.64 |
| Light responsive (n=18) | 49.59 ± 6.88 | 188.28 ± 24.5 | 26.06± 3.9 | -5.21 ± 3.44 |
| **t-value (p-value)** | **-1.76 (0.1)** | **-0.73 (0.47)** | **0.43 (0.67)** | **-0.77 (0.46)** |

**Supplemental Table S1. Passive membrane properties of PnC GlyT2 cells.** Statistical analysis showed no significant differences in the access resistance ($R_a$), membrane resistance ($R_m$), membrane capacitance ($C_m$), time constant ($\tau$) and holding current ($I_h$) of light responsive and unresponsive $GlyT2^+$ PnC neurons. Data are from n=18 $GlyT2^+$ cells responsive to light and n=20 $GlyT2^+$ cells unresponsive to the photostimulation.
